# iBrAVE: a unified framework for 3D interactive and integrative analysis of brain atlas data across modalities and scales

**DOI:** 10.1101/2025.10.04.680445

**Authors:** Yu-Fan Wang, Ming-Quan Chen, Jia-Fei Wei, Hong-Bo Ma, Yong-Wei Zhong, Fu-Ning Li, Chen-Xi Jin, Si-Xuan Cheng, Jia-Qi Guan, Yuan-Ming Zheng, Tian-Lei Zhang, Chao Chen, Xiu-Dan Zheng, Sheng-Jin Xu, Kai Wang, Jiu-Lin Du, Xu-Fei Du

**Author notes:** These authors jointly supervised this work: Jiu-Lin Du, Xu-Fei Du.

## Abstract

Understanding the brain requires integrating molecular, structural, and functional information across multiple scales. Despite the rapid expansion of multimodal, multiscale brain atlases in various species, integrative analysis remains challenging. Here, we present iBrAVE, an open-source platform for 3D interactive, integrative analysis of brain atlas datasets across modalities and scales within a common reference space. iBrAVE unifies heterogeneous datasets into five core representations, enabling source-independent analyses from insects to primates. At the single-modality level, it features spatially resolved quantification, as exemplified by single-neuron morphology analyses spanning geometry, topology, projection, clustering, and putative wiring. Across modalities, it links gene expression and neuronal activity with morphologically defined neurons and putative circuits via establishing spatial correspondence. Across scales, it enables bidirectional inference of neuronal identity and synaptic connectivity between light- and electron-microscopy reconstructions via neuronal morphology matching. Thus, iBrAVE provides a unified framework for integrative brain atlas analyses, facilitating data-driven discovery in neuroscience.

## INTRODUCTION

A comprehensive understanding of the brain requires a holistic view integrating its molecular, structural, and functional dimensions.^1–3^ These distinct types of information need to be presented and integrated within a common reference space to enable cross-modal analysis and interpretation^4^. For example, spatiotemporal patterns of brain-wide neuronal activity can be understood when contextualized by the underlying anatomical connectivity among regions and constituent neurons. Technological advances have enabled the generation of multimodal brain atlas datasets at multiple spatial scales, increasingly spanning developmental stages and diverse species.^5–28^ Unlocking the potential of these large-scale efforts requires tools that integrate, interrogate, and analyze multimodal, multiscale data within a coherent three-dimensional (3D) spatial reference.^29^

High-resolution 3D reference brain atlases have been established for major model organisms,^11,12,15,30–36^ offering standardized anatomical coordinate frameworks for spatial alignment of diverse datasets from micro- to macroscopic scales. Complementary advances in image registration pipelines have further accelerated atlas-based data integration.^37–41^ Building on these foundations, a growing ecosystem of atlas-centered web or software platforms has emerged. Initial web-portal efforts for mouse data emphasized specific brain organizations, such as cell-type cytoarchitecture in the Blue Brain Cell Atlas^42^ and single-neuron reconstructions in the MouseLight NeuronBrowser.^18^ In parallel, web-based portals were developed to provide community access to atlas datasets in other model systems, such as mapzebrain for zebrafish^12^ and Virtual Fly Brain for *Drosophila*.^43^ Alongside these species-specific portals, standalone tools were developed to address common analytical needs: NeuTube supports visualization of SWC neuronal reconstructions,^44^ WholeBrain enables segmentation, registration, and neuroanatomical analysis of light-microscopy images,^45^ and the natverse toolkit provides quantification and clustering of neuronal morphologies.^46^ Moving forward, Brainrender^47^ enables 3D rendering of multiple atlas-registered data, including regions, points, and neuronal tracings. Locare^48^ introduces a standardized workflow to represent heterogeneous neuroscience data as geometric objects in 3D atlases, enabling co-visualization of multimodal datasets within a shared coordinate framework.

These efforts together illustrate a transition from specialized tools tailored to single modalities or species toward frameworks that can accommodate datasets across modalities and species, and from static data exploration to spatially resolved analyses. Despite these advances, a major challenge remains: developing platforms that enable integrative analyses of brain atlas datasets across modalities, scales, and species, thereby supporting holistic understanding of brain structure and function.^29^

To address this issue, we developed the Integrated Brain-data Analysis and Visualization Engine (iBrAVE), an open-source software platform for 3D interactive and integrative analysis of brain atlas data across modalities and scales. iBrAVE supports five core data types: volumetric images, parcellated regions, spatial feature points, neuronal tracings, and networks. These data types collectively provide a comprehensive framework for representing the molecular, structural, and functional dimensions of brain atlas data across micro-, meso-, and macroscopic scales. For each type, iBrAVE offers spatially resolved quantification with dedicated 3D visualization. Once datasets are registered to a common reference atlas, iBrAVE enables cross-modal and cross-scale integrative analyses based on spatial correspondence, regardless of origin and format. Within a unified interface, users can interactively link gene expression patterns and brain-wide neuronal activities with morphologically reconstructed neurons, allowing molecular and functional annotations of targeted neurons and putative circuits. Moreover, it can match light microscopy (LM) and electron microscopy (EM) reconstructions, enabling transfer of neuronal identity from LM to EM data and synaptic connectivity from EM to LM data.

To extend applicability, iBrAVE includes a registration module to align unregistered datasets to species-specific reference templates. In addition, users can incorporate custom algorithms and adapt the platform to emerging data types or analytical needs. To facilitate usability, the platform features a context-sensitive right-click menu, enabling direct interaction with data objects and streamlined access to core visualization and analytical functions. To support computationally intensive tasks, the platform is powered by a high-performance backend infrastructure that ensures real-time rendering and smooth interaction with large-scale datasets, such as whole-brain neuron reconstructions and synaptic-resolution connectomes. Moreover, iBrAVE facilitates data sharing and reproducibility through standardized metadata encoding (JSON) and exportable analytical outputs in Excel format. It also supports high-quality export of static images and user-defined animations for effective scientific communication.

Collectively, iBrAVE offers an all-in-one solution for interactive and integrative analysis of brain atlas data across modalities and scales in diverse species. By bridging molecular, structural, and functional domains through spatial contexts, iBrAVE provides an essential foundation for data-driven discovery in neuroscience research.

## RESULTS

### System architecture of the iBrAVE platform

Mapping brain atlas efforts have generated increasingly diverse, large-scale datasets across experimental paradigms, data modalities, and species. These resources often remain fragmented as isolated data silos, and effective use of them is restricted to individual research groups with advanced computational expertise. In developing iBrAVE, we aimed to address these barriers through two guiding design principles. First, we sought to enable spatially grounded integration across diverse datasets registered into a common coordinate space, thereby breaking inter-dataset isolation and fostering cross-modal correspondence. Second, we prioritized accessibility, ensuring that experimenters, regardless of programming experience, could intuitively explore, analyze, and interpret complex 3D brain datasets. To achieve these, we evaluated existing open-source solutions and selected ParaView^49^ as a prototype system, given its strengths in 3D rendering, spatial queries, and scalable data handling. iBrAVE extends this architecture with neuroscience-specific functionalities to support single-modal, cross-modal, and cross-scale brain atlases’ interactive visualized analysis and integration (Figure 1).

**Figure 1.**
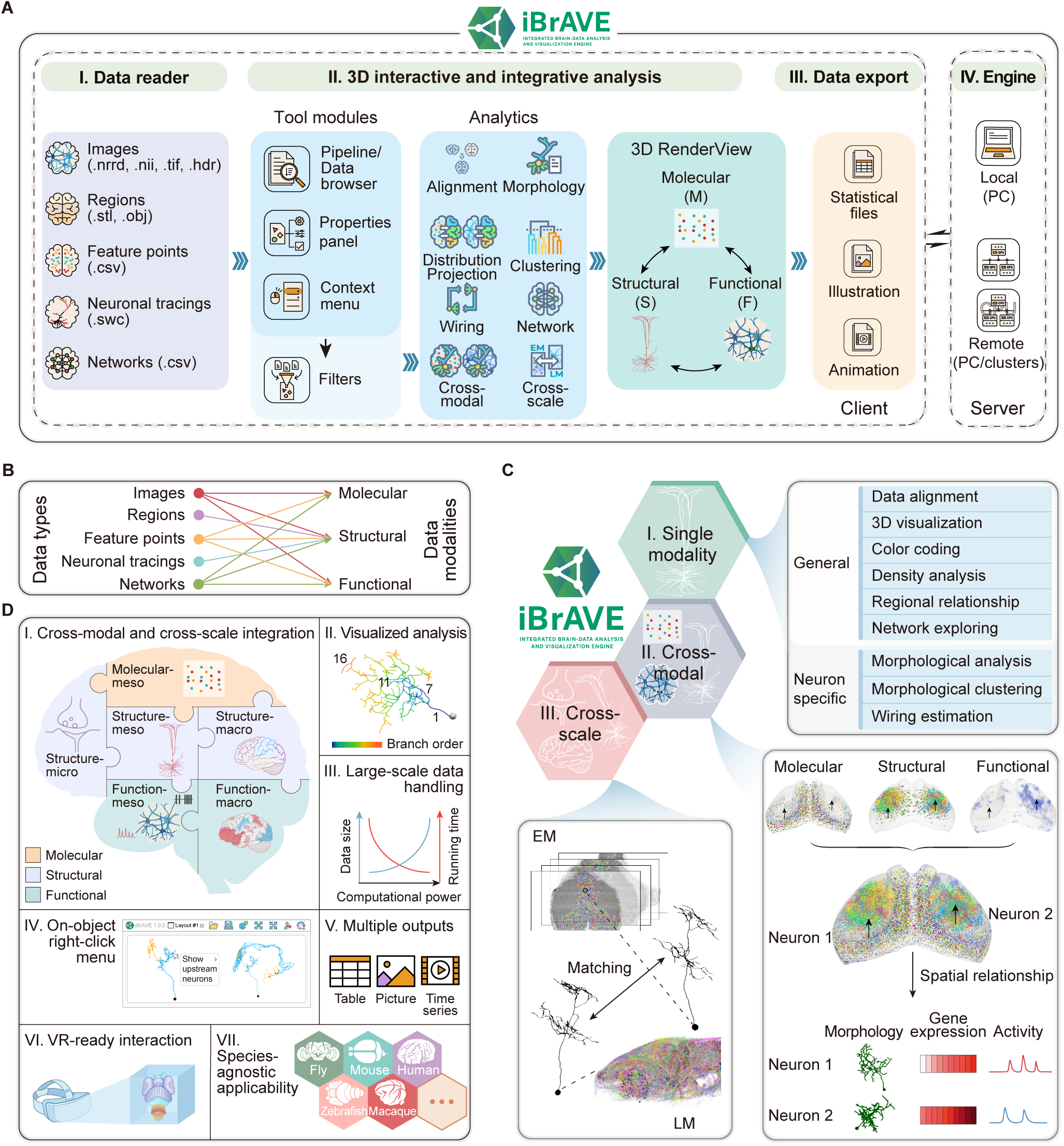
Overview of iBrAVE: system architecture, core functions and key features. (A) System architecture and workflow. iBrAVE comprises a client interface (I–III) and a server engine (IV) that can run locally or on a remote high-performance cluster. The client manages multimodal data import (I), processes it via five tool modules that collaboratively enable multiple analytic functions mediated by the *Filter* (II, left), and performs single-modality and multimodal integrative analyses in an interactive 3D viewer (II, right). Results can be exported as tables, pictures, or videos (III). (B) Correspondence between supported data types (left) and data modalities (right). (C) Core functions. (I) Single-modality analysis (top-right), offering both general and modality-specific tools. (II) Cross-modal integrative analysis (bottom-right), enabling spatial mapping of molecular, structural, and functional data at cellular resolution to assign multimodal annotations to individual neurons. Arrows mark the soma position of two representative neurons. (III) Cross-scale matching (bottom-left), allowing bidirectional correspondence between electron microscopy (EM) and light microscopy (LM) reconstructions. (D) Key features. (I) Integration of multimodal datasets across spatial scales. Background colors denote data modalities. (II) Visualization of quantitative analyses, as exemplified by coloring neuronal morphology according to neurite branch order. Numbers indicate the order of four representative branches. (III) Large-scale data handling. The schematic shows that, as computational power increases, data size (blue) increases while running time (red) decreases. (IV) Interactive 3D object-based queries via a context menu. As exemplified, right-clicking on an optic tectum (OT) neuron (left) allows retrieve of its presynaptic retinal ganglion cells (RGCs) (right). Soma, black; dendrite, yellow; axon, cyan. (V) Flexible export of results, including tables, pictures, and videos. (VI) Immersive data exploration in virtual reality (VR). (VII) Species-agnostic compatibility, supporting brain atlases of fly, zebrafish, mouse, macaque, human, and other model organisms. See also Videos S1 and S5.

The first step was to standardize the import and representation of heterogeneous datasets across species. Despite methodological diversity, most brain atlas data fall into three principal modalities—molecular, structural, and functional—each represented at microscopic (i.e., synaptic level), mesoscopic (i.e., single-cell or cell-type level), and macroscopic (i.e., brain-region level) scales. To accommodate this diversity, iBrAVE was designed to ingest five widely applicable data types (Figures 1A-I and 1B): (1) volumetric images (.nrrd, .nii, .tif, .hdr), the primary representation of raw data; (2) parcellated regions (.stl, .obj), 3D surfaces delineating brain regions or structures; (3) spatial feature points (.csv), positions of discrete biological features, such as gene expression signals, labeled proteins or cells, axon terminals, and functionally active neurons; (4) neuronal tracings (.swc), skeletonized reconstructions of individual neurons from LM and EM datasets; and (5) networks (.csv), graph-based source-target matrices representing interactions among biological entities, including neurons, neuronal types, or brain regions. Please note that .stl, .obj, .swc, and .csv files are digitized data derived from image datasets. This species-agnostic, source-independent framework enables integration of datasets spanning spatial transcriptomics, transgenic labeling, neuronal reconstructions from LM and EM, functional recordings such as calcium imaging or functional magnetic resonance imaging (fMRI), and multiscale network maps.

Once loaded, datasets can be explored and analyzed through a modular graphical interface comprising multiple coordinated tools (Figure 1A-II; see Methods). The *Pipeline Browser* and *Data Browser* serve complementary roles in data management. The *Pipeline Browser* organizes the processing workflow, enabling users to launch analysis modules (*Filters*) and adjust their parameters via the *Properties Panel*. The *Data Browser* focuses on the dataset itself, providing control over the visibility, sorting, and coloring of individual elements. The 3D *RenderView* provides spatially resolved visualization and direct user interaction (Figure 1A-II). Quantitative analysis results are rendered within this view, for example, neuronal morphologies are colored according to branch order (Figure 1D-II). We created a *Context Menu* within the 3D *RenderView* to facilitate rapid toggling of object visibility, color customization, grouping, and export of analysis results. The 3D *RenderView* supports spatially driven interactions with the data: users can select objects directly in 3D space and dynamically retrieve mapped information, such as displaying presynaptic neurons of a chosen neuron (Figure 1D-IV). The entire process is fully visualized, enabling transparent and intuitive data interrogation.

The analytical functions of iBrAVE are implemented through the *Filters* module (Figure 1A-II), which can be directly selected to apply or triggered via the *Context Menu*. These functions fall into three hierarchical categories (Figure 1C): (1) single-modal quantification (Figure 1C, top right; see also Figure 3), including general analysis (e.g., density and regional distribution) and modality-specific analysis (e.g., for the neuronal structural data, iBrAVE can calculate morphological parameters and similarity, and estimate putative connections between individual neurons based on axon-dendrite proximity); (2) cross-modal mapping between molecular, structural, and functional data, linking gene expression profiles, neuronal morphology, and neural activity patterns based on spatial queries (Figure 1C, bottom right; see also Figures 4 and 5); (3) cross-scale integration (Figure 1C, bottom left; see also Figure 6), matching single-cell reconstructions across LM and EM data to bridge microscopic connectivity with mesoscopic genetic and functional features. Collectively, these functions achieve integration across modalities and scales (Figure 1D-I). For unaligned datasets, the Alignment in the *Filter* provides template-based registration (powered by ANTs) of images and their derived digitized data.

**Figure 2.**
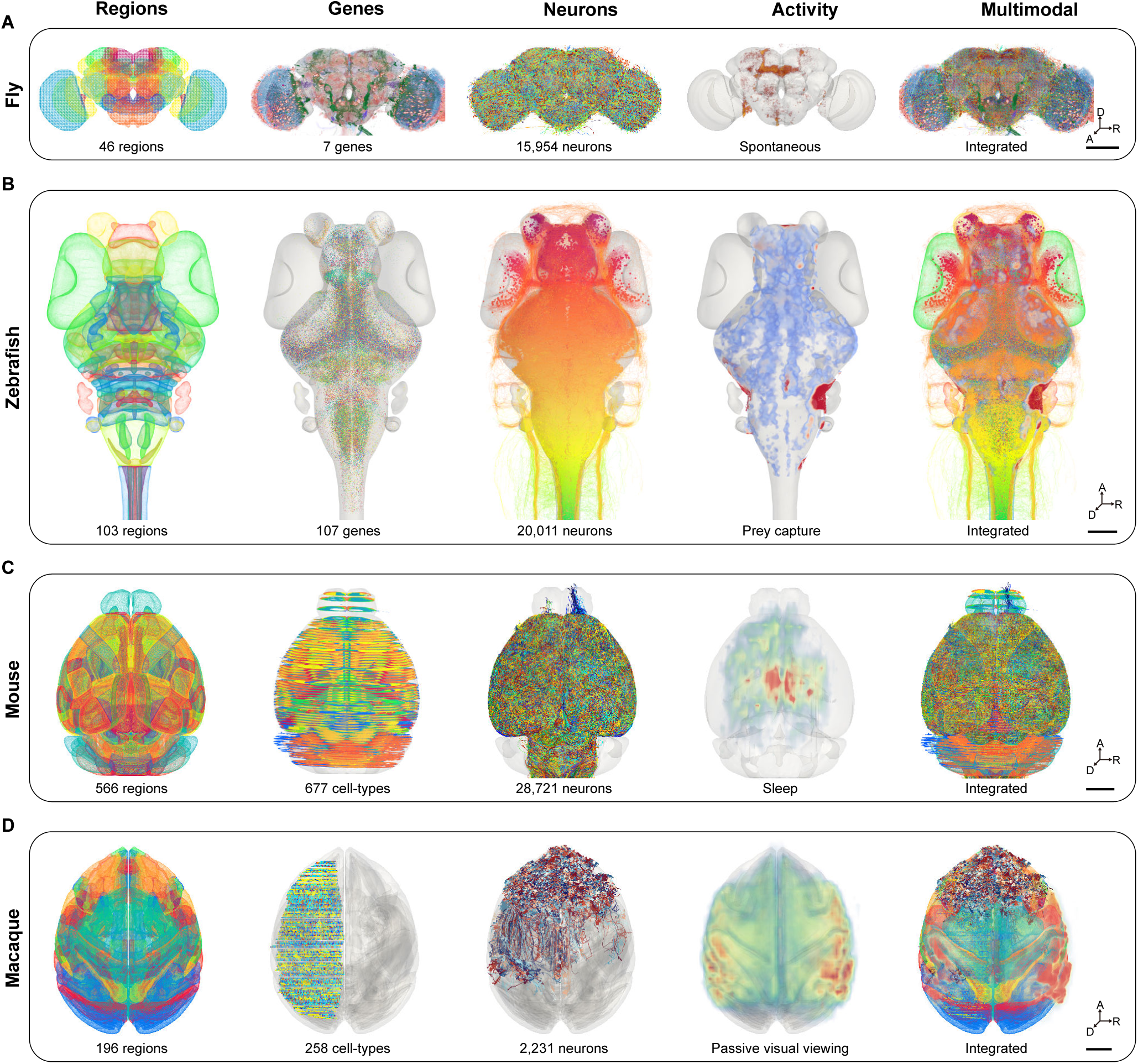
Co-visualization of multimodal data in 3D reference atlas across species. Left to right: anatomical regions, spatial gene expression, neuronal tracings, functional activity maps, and multimodal overlay for fly (A), zebrafish (B), mouse (C), and macaque (D). (A) Fly brain reference atlas (JRC 2018) with 46 regions,^30^ 7 gene-expression images,^50^ and 15,954 single-neuron morphology tracings,^51^ all color-coded by identity; brain-wide spontaneous neuronal calcium activity,^53^ color-coded by fluorescence intensity; integrated cross-modal visualization of gene expression, neuronal tracing, and neuronal activity datasets. (B) Zebrafish brain reference atlas (ZExplorer CPSv1.0) with 103 regions,^15^ 107 gene-expression maps,^14^ and 20,011 single-neuron morphology tracings,^15^ all color-coded by identity; brain-wide neuronal calcium activity during the approach and eye convergence epoch of prey capture behavior (see Figure 5), color-coded by ΔF/F_0_; integrated cross-modal visualization of gene expression, neuronal tracing, and neuronal activity datasets. (C) Mouse brain reference atlas (Allen CCFv3) with 566 regions,^32^ 677 cell-types inferred from spatial transcriptomics,^22^ and 28,721 single-neuron morphology tracings (18,621 from prefrontal cortex^20,24^ and 10,100 from hippocampus^23^), all color coded by identity; fMRI activity during sleep,^54^ color-coded by image intensity; integrated cross-modal visualization of gene expression, neuronal tracing, and fMRI activity datasets. (D) Macaque brain reference atlas (NMTv2) with 196 regions,^36^ 258 cell-types inferred from spatial transcriptomics,^26^ and 2,231 single-neuron morphology tracings,^27^ all color coded by identity; fMRI activity during passive visual viewing,^55^ color-coded by image intensity; integrated cross-modal visualization of gene expression, neuronal tracing, and fMRI activity datasets. Scale bars: 100 μm (A and B), 2 mm (C), 10 mm (D). A, anterior; D, dorsal; R, right. See also Figure S1, Tables S1 and S2, and Video S1.

**Figure 3.**
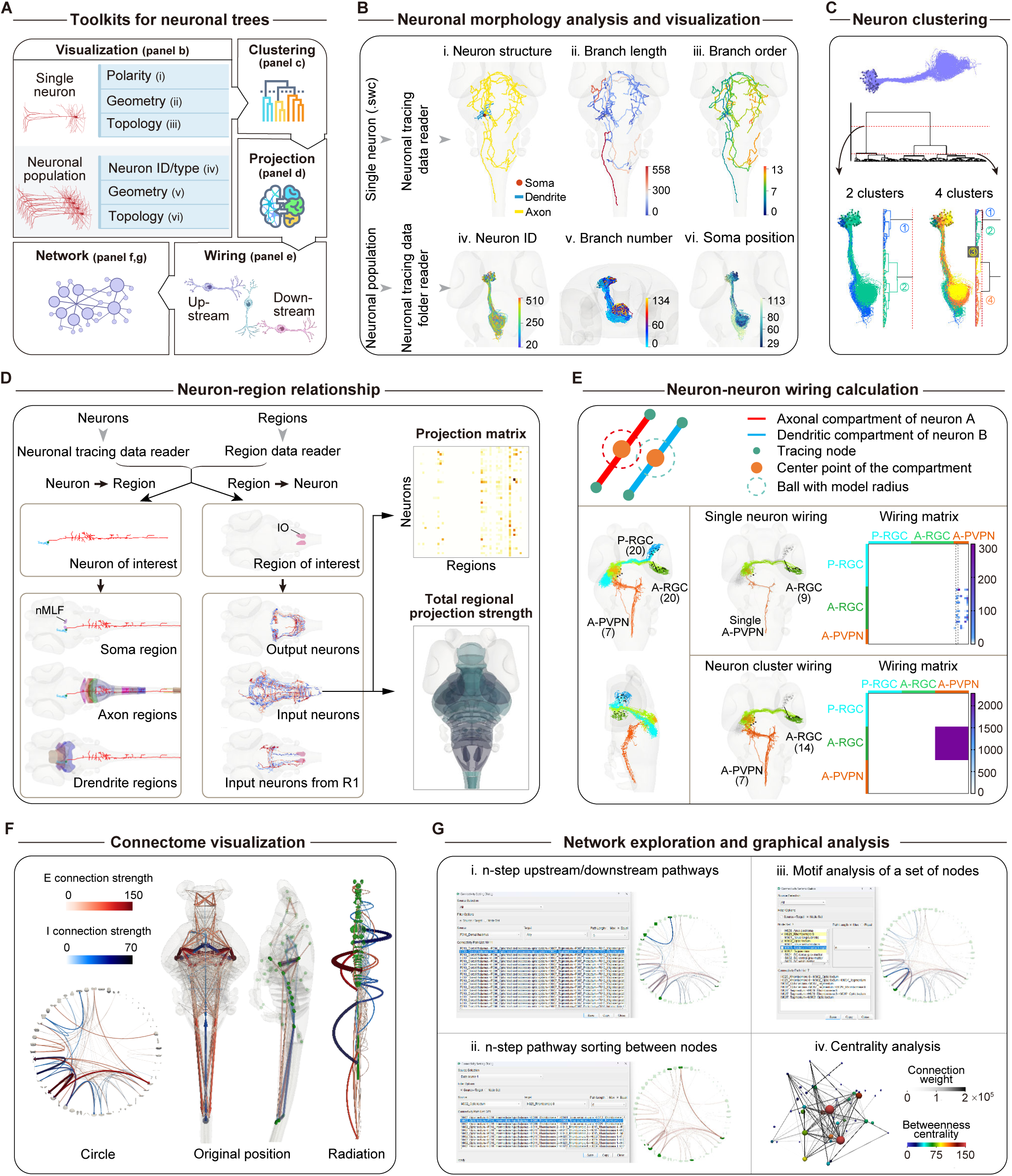
Analytical toolkits for neuronal structures. (A) Toolkits for neuronal structure analysis comprising modules for visualization, clustering, projection mapping, wiring calculation, and network analysis. (B) Neuronal morphology analysis and visualization. Top: single-neuron morphology renderings, color-coded by morphological compartments (i), branch length (ii), or branch order (iii). Bottom: population-level visualization with neurons color-coded by identity (iv), branch number (v), or somatic position (vi). (C) Interactive clustering of neuronal populations. Input neurons (top) with dendrogram (middle) cut at two levels (red dashed lines), yielding either two clusters (bottom left) or four clusters (bottom right). Neurons and dendrogram branches are color-coded by cluster identity. (D) Neuron-region relationship exploration. Left (neuron-to-region query): an nMLF neuron showing soma location, axonal projection targets, and dendritic innervation regions. Middle (region-to-neuron query): neurons located in (output) and projecting to (input) the IO, or projecting to the IO from the specific region R1. Right (projection strength analysis): single-neuron projection matrix (top) and regional projection map (bottom, color-coded by total projection strength) of IO input neurons. The projection strength is calculated as total axonal length terminating in each region. (E) Neuron-neuron wiring analysis. Top: schematic of the ‘center’ computational mode for inferring connectivity by axodendritic spatial proximity (see Methods). Bottom left: RGCs targeting anterior (green) or posterior (blue) optic tectal neuropil (A-RGCs and P-RGCs), and a PVPN subtype (orange, A-PVPN) included in analysis. Bottom right: circuit rendering (left, with all RGCs and PVPNs shown in grey as background) and wiring matrices (right) at single-neuron (top) and cluster (bottom) levels. The connection strength is quantified as the number of axon-dendrite compartment pairs within a 1-μm radius. The area between the two dashed lines marks the upstream RGC connectivity to the single A-PVPN. Numbers in brackets indicate the number of neurons visualized. (F) Putative connectivity visualization. Zebrafish interregional excitatory (red) and inhibitory (blue) connections^15^ rendered in multiple layouts: circular (regions arranged in a ring, left), 3D anatomical (without or with region-centroid spheres, middle), and radial (regions arranged by *y*-axis position, right). The connection strength is encoded by edge color and width, and additionally by parabola height in the radial layout. (G) Network analysis toolkit. Left: network exploration of the connectome in (F), including (i) six-step downstream pathways from the dorsal thalamus and (ii) six-step paths between the OT and R8. Right: graph-theoretic analyses, including (iii) motif detection within selected node sets (R8, OT, and tegmentum) and (iv) centrality analysis, with randomly arranged nodes sized and colored by betweenness centrality, and edges brightness-coded by connection weight. For i–iii, the selected pathways highlighted in a circular layout (right). IO, inferior olive; nMLF, nucleus of the medial longitudinal fasciculus; OT, optic tectum; PVPN, periventricular projection neuron; R1/8, rhombomere 1/8; RGC, retinal ganglion cell. See also Videos S2 and S3.

**Figure 4.**
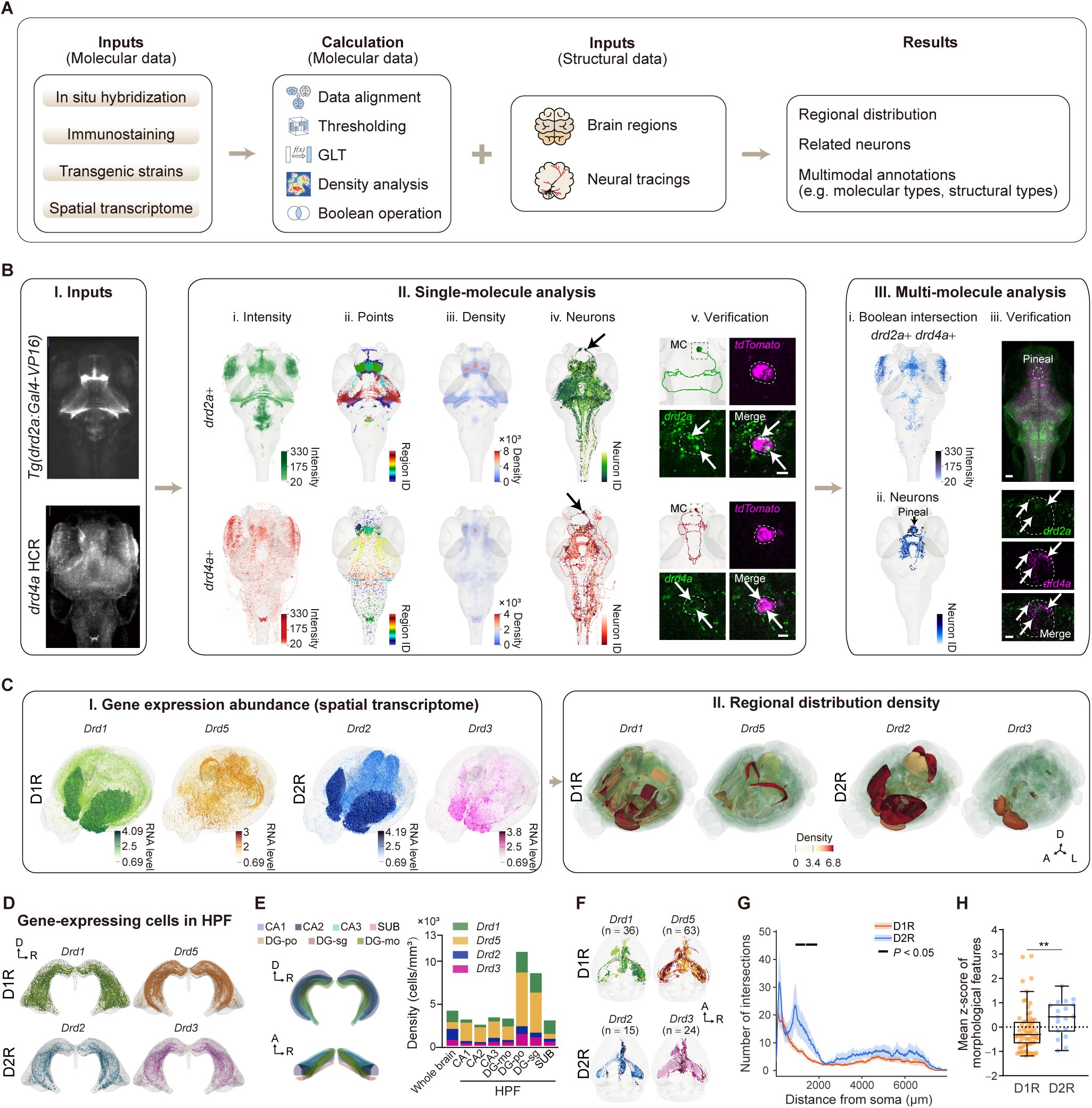
Mapping gene expression to neuronal morphology. (A) Workflow of gene-to-structure analysis. Registered gene expression images and spatial point data undergo various computational processes. Raw or intermediate outputs can be mapped onto structural datasets to reconstruct regional gene expression, identify gene-expression-related neurons, and generate multimodal annotation. (B) Demonstration of gene-to-structure mapping using gene expression images in zebrafish. (I) Example raw images: *Tg(drd2a:GAL4-VP16)* fluorescence (top, crossed to *Tg(UAS:Dendra-kras)*) and *drd4a* HCR (bottom). (II) Single-molecule analysis: visualization as a 3D intensity image (i); digitization into spatial feature points, color-coded by regional distribution (ii) and density (iii); spatial mapping onto single-neuron morphology reconstructions^15^ to filter candidate neuronal tracings (iv), including two mitral cells (MCs, arrow); experimental validation of gene expression by HCR-FISH on the two MCs (v), which expressed tdTomato for morphology tracing (see Methods). Green puncta, HCR-FISH signals; white arrows, co-localization of HCR-FISH signals and tdTomato; dashed circles, outlines of MC somata. (III) Multi-molecule analysis: (i) Boolean intersection of the *drd2a* and *drd4a* expression images; (ii) reconstructed pineal gland neurons^15^ (arrow) identified by spatial mapping to the intersection pattern; (iii) HCR-FISH revealed the juxtaposition of *drd2a*^+^ (green) and *drd4a*^+^ (magenta) puncta within a single optical plane in the pineal gland (dashed circle), supporting co-expression of *drd2a* and *drd4a* in pineal neurons. White arrows indicate co-expression puncta. Scale bars: 5 μm (II-v), 50 μm (III-iii, top), 10 μm (III-iii, bottom). (C–F) Demonstration of gene-to-structure mapping in mice. (C) Mouse spatial transcriptomics visualized in iBrAVE. (I) Expression abundance (normalized RNA counts) of dopamine receptor genes (D1R: *Drd1* and *Drd5*, D2R: *Drd2* and *Drd3*) at single-cell resolution. (II) Region-wise density of neurons expressing the four genes (×10^3^ cells per mm^3^). (D) Spatial distribution of *Drd1*-, *Drd5*-, *Drd2*-, and *Drd3-*expressing neurons in the HPF. (E) Subregional organization of the HPF (left) and densities of *Drd1*-, *Drd5*-, *Drd2*-, and *Drd3-*expressing neurons in HPF subregions (right). (F) Volume rendering of the corresponding HPF neurons in the single-neuron morphology reconstruction dataset^23^. These neurons were identified by mapping *Drd1*, *Drd5*, *Drd2*, and *Drd3* expression patterns onto the morphology reconstruction dataset through the spatial correspondence function in iBrAVE. The number in the brackets indicates the number of reconstructed neurons identified. (G) 3D Sholl analysis of neurons exclusively mapped to D1R (n = 65) or D2R (n = 18). Intersections were quantified at 10-μm radial intervals from the soma center using concentric spheres. Solid lines represent mean values, and shaded areas indicate SEM. Horizontal black lines above the plots indicate individual radii with significant difference (*P* < 0.05; Wilcoxon rank-sum test with false discovery rate correction). (H) Box-and-whisker plots with overlaid individual data points showing the mean z-score of morphological features for neurons exclusively mapped to D1R (n = 65) or D2R (n = 18). The score was calculated for each neuron as the mean z-score across five morphological features: total length, bifurcation number, branch number, terminal number, and maximum branch order. The plots were generated in Tukey style, where the center line indicates the median, the box spans the interquartile range (IQR; 25th - 75th percentiles), and whiskers extend to the most extreme data points within 1.5 × IQR. ***\*\*****P* < 0.01 (Wilcoxon rank-sum test). A, anterior; D, dorsal; L, left; R, right. D1/2R, D1/2 dopamine receptor; HPF, hippocampal formation; GLT, gray level transformation.

**Figure 5.**
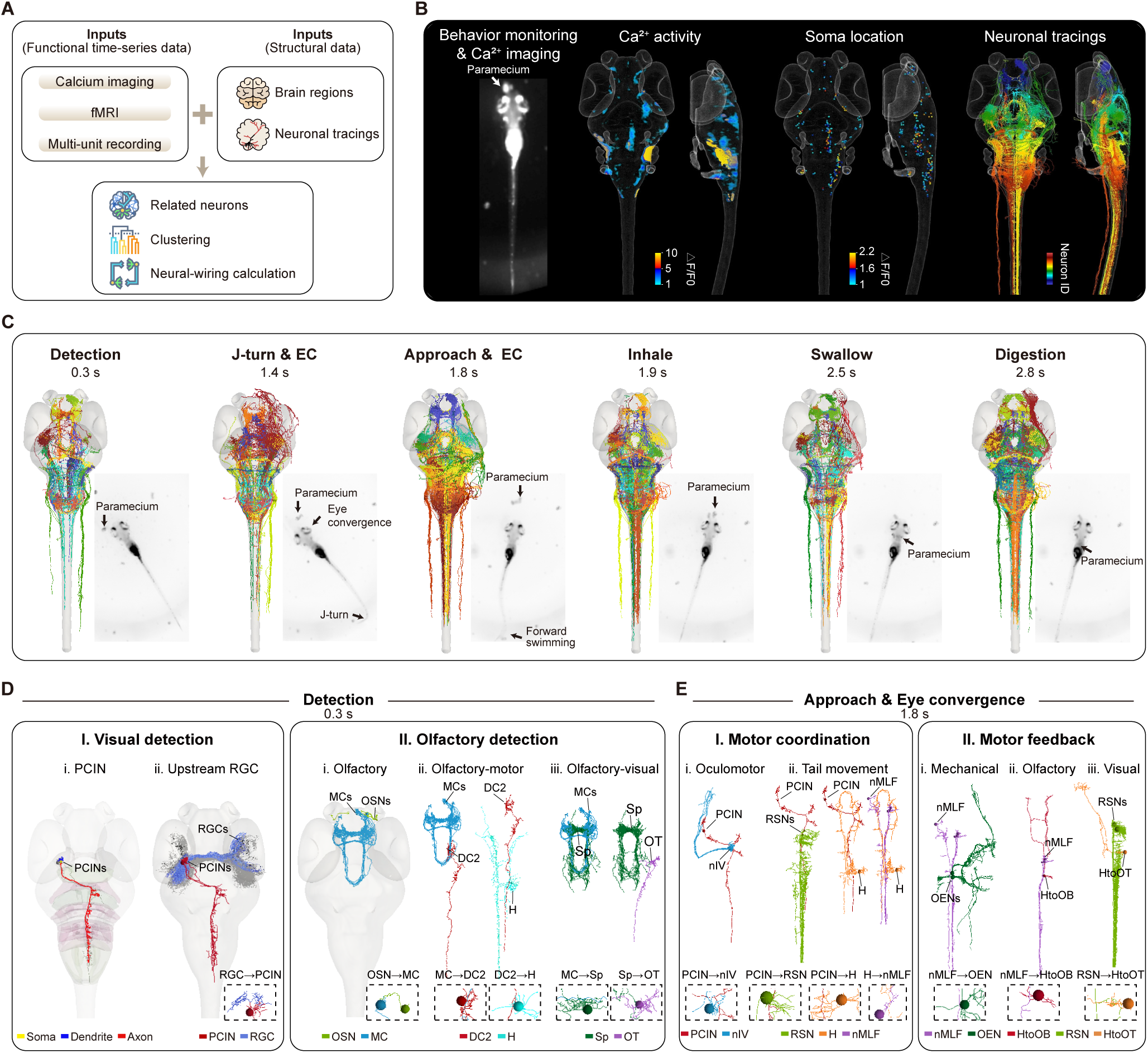
Mapping whole-brain neuronal activities to neural circuits. (A) Workflow of function-to-structure analysis. Registered functional imaging time-series (e.g., calcium imaging, fMRI) or time-series activity matrices of cells with spatial coordinates (e.g., multi-unit recording) can be imported into iBrAVE to map corresponding neuronal tracings, cluster mapped neurons, and identify potential neural circuits. (B) Alignment of behavior, neuronal activity, and single-neuron morphology tracings of larval zebrafish. Using the spatial correspondence function in iBrAVE, brain-wide neuronal Ca^2+^ activity (second on the left, color coded by ΔF/F_0_) during prey capture (first on the left) was in real-time mapped to the soma positions (third on the left, color-coded by ΔF/F_0_ at each soma) and morphologies (right, color-coded by anteroposterior soma position) of corresponding neurons in the single-neuron morphology reconstruction dataset.^15^ (C) Architecture of potential circuits during different prey-capture epochs. One time point across each of six behavioral epochs (detection, J-turn & eye convergence (EC), approach & EC, inhale, swallow, and digestion) is shown with the morphologies of corresponding reconstructed neurons, which are color-coded by excitatory and inhibitory morphotypes.^15^ Bottom right, corresponding larval behavior and paramecium position. (D and E) Potential circuit dissection of prey detection (D) and approach & EC (E) epochs at the time point in (C), using neuronal wiring analysis. For prey detection, corresponding neurons include both visual (D-I, RGCs→PCINs) and olfactory (D-II-i, OSNs→MCs) pathways. Downstream of MCs, corresponding neurons engage motor (D-II-ii, spinal cord-projecting DC2 and hindbrain neurons) and visual (D-II-iii, OT neuron via Sp) circuits. For approach & EC, downstream of PCIN, corresponding neurons include motor-related neurons for oculomotor (E-I-i, nIV) and tail movement (E-I-ii, RSNs and nMLF). Further downstream of RSNs and nMLF are hindbrain neurons projecting ascendingly to mechanosensory (E-II-i, OEN), olfactory (E-II-ii, HtoOB), and visual (E-II-iii, HtoOT) systems. Neuronal tracings are color-coded by morphotypes^15^. Dashed boxes highlight representative contacts between upstream and downstream morphotypes. DC2, diencephalic cluster 2; H, hindbrain; HtoOB, hindbrain-to-OB projecting neurons; HtoOT, hindbrain-to-OT projecting neurons; MC, mitral cell; nMLF, nucleus of the medial longitudinal fasciculus; nIV, trochlear nucleus; OEN, octavolateral efferent nucleus; OSN, olfactory sensory neuron; OT, optic tectum; PCIN, prey-capture initiation neuron; RGC, retinal ganglion cell; RSN, reticulospinal neuron; Sp, subpallium. See also Video S4.

**Figure 6.**
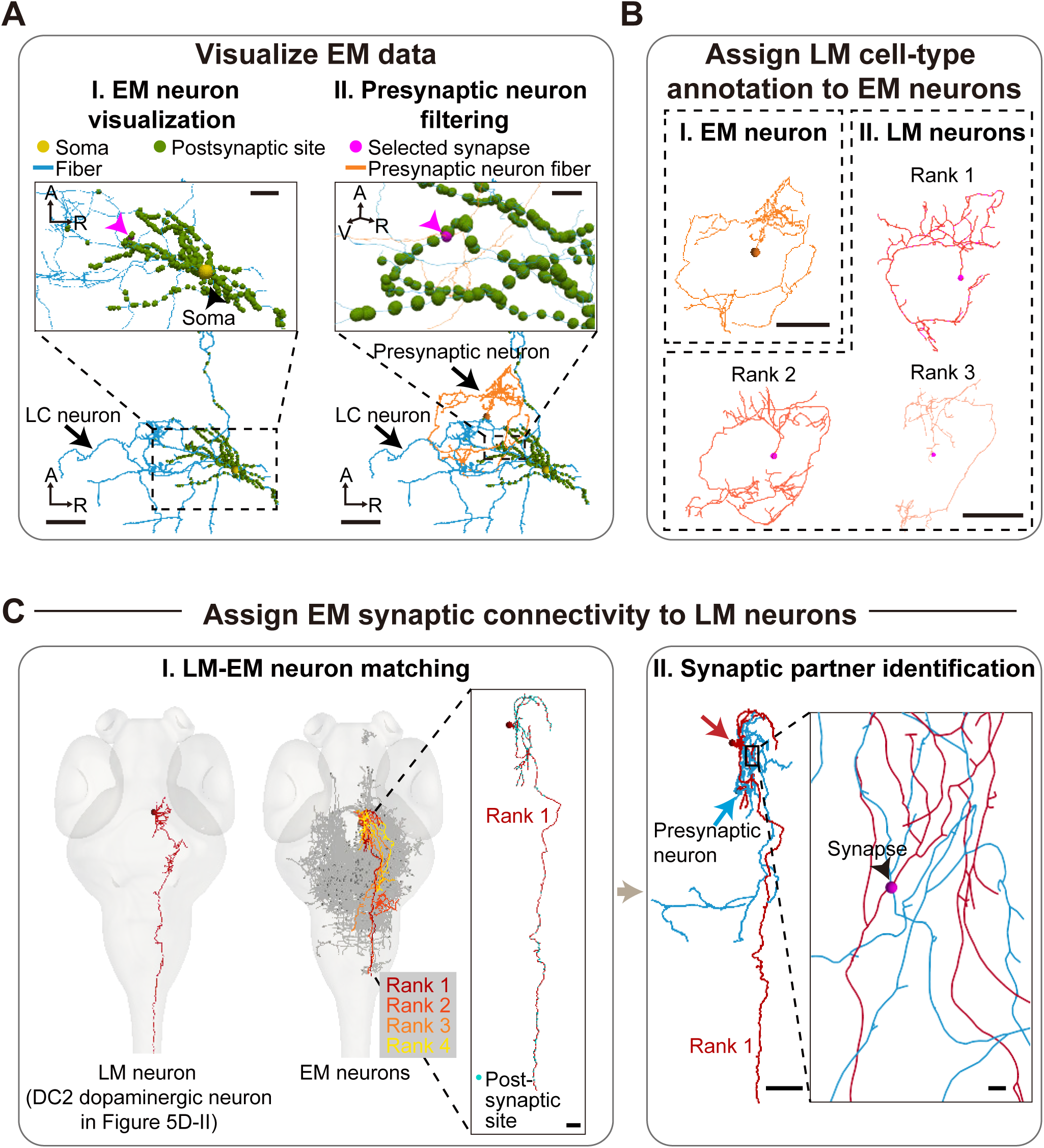
Bidirectional linking of LM and EM datasets through morphological matching. (A) Volume rending of an EM-reconstructed locus coeruleus (LC) neuron (yellow, soma; cyan, neurite; green, postsynaptic sites) (I), and co-rendering with a presynaptic partner (orange) filtered by a postsynaptic site (magenta arrowhead) (II). The black box in (II) shows a rotated and enlarged view of the inset (dashed rectangle), illustrating the selected postsynaptic site. (B) Filtered presynaptic neuron in (A) (I) and the three-highest scoring morphological matches from the LM structural atlas^15^ (II). These LM neurons belong to a *vglut2a/2b*+ morphotype (# Glu-5L0701^15^) located in the dopaminergic diencephalic cluster 6. (C) Functionally defined prey capture-activated LM dopaminergic neuron in DC2 (I, left; see Figure 5D, II-ii) and matched neurons (I, middle, color tracings) from the EM atlas (I, middle, grey tracings^16^). The top-matched EM neuron (rank1) is enlarged with postsynaptic sites in cyan (I, right). A presynaptic partner (cyan) of this rank1 EM neuron (red) was further identified, forming a single synapse (arrowhead) between them (II). Arrows indicate the somata of the two neurons. Scale bars: 50 μm (A-I,II, B, C-II), 20 μm (A-I, inset), 5 μm (A-II, inset; C-I,II, inset). A, anterior; R, right; V, ventral. DC2, diencephalic cluster 2; EM, electron microscopy; LM, light microscopy.

At the output end, iBrAVE supports flexible export options ranging from quantitative tables for downstream analysis to static figures and spatiotemporal animations (Figures 1A-III and 1D-V). A client-server architecture (Figure 1A-IV) ensures scalability and high-performance visualization and analysis of large-scale datasets (Figure 1D-III). All visualizations are compatible with immersive exploration in virtual reality (VR) for enhanced spatial perception and interaction (Figure 1D-VI). These features make the platform both intuitive and practical.

Taken together, iBrAVE supports an end-to-end workflow from multimodal data ingestion and spatial registration to quantitative, cross-modal, and spatially resolved integrative analysis, all within a unified, interactive, and scalable 3D environment, without species restrictions (Figure 1D-VII).

### Data handling capability for multimodal atlas datasets across species

To verify that iBrAVE provides a unified environment for ingesting and visualizing multimodal brain atlas datasets across species, we curated publicly available resources spanning the three major modalities—molecular (gene expression), structural (single-neuron morphology), and functional (brain-wide activity)—from four widely studied model organisms ranging from invertebrates to non-human primates: *Drosophila melanogaster* (fly), *Danio rerio* (zebrafish), *Mus musculus* (mouse), and *Macaca mulatta* (macaque) (Figure 2). For fly, we adopted the JRC 2018 central brain template^30^ (Figure 2A, left). For zebrafish, we used our nervous system-wide 3D Common Physical Space (CPSv1)^15^ as the reference atlas for integrating multimodal datasets (Figure 2B, left). For mouse and macaque, we adopted the Allen CCFv3^32^ (Figure 2C, left) and the National Institute of Mental Health Macaque Template NMTv2^36^ (Figure 2D, left), respectively.

Molecular datasets, in the form of registered transgenic reporter or *in situ* hybridization images, can be directly loaded into iBrAVE and co-visualized with brain structures (Figure 2A, second column from the left, gene expression images of fly^50^ and Table S2). To enable cell-level visualization, we converted gene expression images into spatial feature point representations, where the 3D coordinates of gene-expressing cells were recorded as individual markers (see Methods). This allows specific gene-labeled cells or transcriptomic cell types to be visualized as color-coded points in 3D. Datasets converted in this way included zebrafish whole-brain hybridization chain reaction (HCR) images^14^ (Figure 2B, second column from the left and Table S2), mouse MERFISH-based spatial transcriptomics^22^ (Figure 2C, second column from the left), and macaque cortical stereo-seq data^26^ (Figure 2D, second column from the left). Structural datasets consisted of large-scale single-neuron reconstructions, which had already been registered to their respective templates of the four species,^15,20,23,24,27,51^ were directly visualized in iBrAVE via standardized SWC^52^ file format (Figures 2A–2D, middle). Functional activity datasets, such as *in vivo* calcium imaging in fly^53^ and zebrafish, or fMRI in mouse^54^ and macaque,^55^ were likewise ingested as volumetric images and registered to the templates for whole-brain visualization (Figure 2A–2D, second column from the right). Thus iBrAVE enables the integrated visualization of molecular, structural, and functional datasets in a common anatomical reference frame (Figure 2A–2D, right and Video S1), providing a foundation for spatial comparison and cross-modal analysis. Detailed information about the metadata, including the dataset size, rendering complexity, and source, is summarized in Table S1.

To benchmark iBrAVE performance and scalability, we evaluated iBrAVE using 5,000 single-neuron morphology reconstructions from zebrafish and mouse brains, and 2,000 from macaque brains. We quantified five core operations (Figure S1): (1) data reading/parsing (i.e., loading SWCs into iBrAVE’s internal data structure), (2) static rendering (i.e., non-interactive 3D visualization), (3) dynamic rendering (i.e., real-time interactive visualization), (4) neuronal morphology analysis, and (5) neuron-to-region projection strength analysis. Benchmarking was conducted under two deployment modes (see Methods): a standalone workstation and a two-node GPU cluster (dual A100 GPUs per node). Processing times scaled with the number of points per SWC, such that zebrafish neurons, which contain fewer points, were loaded and analyzed faster than mouse neurons despite equal file counts (Figures S1A and S1B). Compared with the standalone workstation, the distributed deployment (cluster) significantly accelerated the operations (Figure S1; average time reduction for all five processes: zebrafish, 82 ± 7%; mouse, 87 ± 3%; macaque, 60 ± 17%). Under the distributed configuration, reading and rendering thousands of neurons required several tens of seconds (Figure S1, sum of the first three operations), while computationally intensive projection analyses were completed within seconds to about ten minutes.

Together, these results show that iBrAVE supports multimodal brain atlas datasets across species, and that distributed environments can enhance its performance.

### Toolkit for neuronal trees: from single-neuron morphology to putative network wiring

Neurons possess distinct dendritic and axonal compartments, which underlie neural connectivity and define the directionality of information flow. To characterize these aspects, iBrAVE implements a modular suite of tools, including structural feature extraction, morphological clustering, analysis of dendritic and axonal projections, inference of potential synaptic partners, and topological network exploration and analysis (Figure 3A).

iBrAVE provides customizable quantitative analysis and visualization for single-neuron morphology reconstructions. Users can apply feature-based coloring schemes to distinguish morphological components such as soma, dendrites, and axons (Figure 3B-i). By integrating core functions of L-measure,^56^ it enables the extraction of key morphological parameters, including total length, Euclidean distance to soma, and branch number and order. These parameters can be exported in Excel table and rendered directly in 3D—for example, coloring each branch by length or order—thus linking geometric and topological features to spatial context (Figures 3B-ii,iii). For neuronal populations, iBrAVE supports batch loading via a data folder and group-wise coloring by neuron ID (Figure 3B-iv), neuronal type (see Methods), individual morphological parameters (such as branch number in Figure 3B-v), or soma position (Figure 3B-vi).

iBrAVE supports neuron clustering based on morphological similarity (Figure 3C). It includes a built-in implementation of MBLAST algorithm^15^ and allows integration of user-defined clustering methods. Users can flexibly define the number of clusters and thus interactively explore the hierarchical organization through correspondingly rendered dendrogram and cluster-specific neuron visualization (Figure 3C, bottom). Neuron clusters can be exported and reloaded for customized visualization and wiring analysis (see below).

iBrAVE integrates projection analysis with real-time 3D interaction, enabling intuitive exploration of neuron-region relationships. Users can directly right-click neurons or brain regions of interest in the *RenderView* to access a context-sensitive menu and invoke analytical visualizations (Video S2). For example, users can select a neuron to query its somatic location, axonal projection regions, and dendritic recipient regions (Figure 3D, left). Conversely, selecting a region reveals its resident neurons (i.e., output neurons) as well as input neurons that project to it, with options for further filtering by the source regions of input neurons (Figure 3D, middle). In addition, users can define custom groups (e.g., excitatory neurons within a nucleus) (see Methods) and perform neuron-region relationship analyses at the group level. Neuron or neuronal cluster projection strength (calculated by axonal length) matrices can be exported as raw tables for downstream analysis or displayed directly in iBrAVE as interactive heatmaps with source-target lookup and connectivity strength visualization (Figure 3D, top right and Video S2). The target regions can also be color-coded to directly display projection strength (Figure 3D, bottom right and Video S2). This integration of interactive querying, computation, and visualization ensures interpretable readouts.

For neuron datasets annotated with axon-dendrite polarity, iBrAVE infers putative synaptic connectivity based on axonal and dendritic spatial proximity within user-defined thresholds (Figure 3E, top; see Methods). This enables reconstruction of microcircuits based on individual neurons or neuronal clusters. For instance, we identified a subtype of periventricular projection neurons (PVPNs) in zebrafish that forms selective connections with retinal ganglion cells (RGCs) targeting the anterior tectal neuropil (Figure 3E, bottom). Inferred connections can be exported (in matrices and JSON format) or interactively queried in the 3D interface by directly picking a specific neuron or neuronal clusters (Figure 3E, bottom middle and bottom right, and Video S3). This interactive connectivity query is applicable to EM-derived neuronal reconstructions.

Beyond upstream-downstream circuit exploration, iBrAVE supports network-level analysis of neuron-, region-, and any node-based graphs, with or without explicit spatial embedding (Figures 3F and 3G). Networks can be visualized as abstract node-edge layouts (i.e., circular), or as spatially grounded views (i.e., original), where nodes are represented by their corresponding anatomical objects or centroids. In addition, regions can be arranged by their centroids, with edges displayed as parabolic arcs (i.e., radiation). Users can specify source-target nodes and path steps to identify all defined communication pathways, which are dynamically linked to 3D network renderings so that selected pathways (including nodes and edges) can be highlighted (Figure 3G-i,ii). Beyond pathway exploration, iBrAVE integrates graph-theoretic analyses to extract higher-order organizational principles. These include motif detection for user-defined node profiles (Figure 3G-iii) and centrality mapping for highlighting communication hubs and key integrator nodes (Figure 3G-iv). This network analysis module can be applied to explore diverse types of connectivity networks written as source-target matrices.

Together, these integrated modules transform raw neuron reconstructions into a versatile analytical substrate, enabling multiscale exploration—from single-neuron morphology to circuit to network topology—within a unified spatial framework. By incorporating wiring inference between individual neurons and graph-based network analysis, iBrAVE extends morphology-centered tools to enable interrogation of how individual neurons assemble into putative circuit architectures.

### Cross-modal analysis: linking gene expression to neuronal morphology

A central objective of iBrAVE is to enable cross-modal integration through spatial correspondence. Structural data are represented primarily by digitized neuronal skeletons (i.e., connected point sets), while molecular and neural activity data are usually expressed as volumetric images, which can be transformed into spatial feature points. Within iBrAVE, raw images can be processed using diverse computational operations, most notably Boolean operations (union, intersection, subtraction), to generate derived images that highlight user-defined features of interest. To enable interaction among these diverse modalities, iBrAVE implements robust spatial query functions across image-image, image-point, and point-point data types. Leveraging the spatial parsing engine, images or feature points can be mapped onto anatomical regions, and particularly neuronal tracings can be directly queried against these data, thereby generating multimodal annotations at the level of individual objects (Figure 4A; see Methods).

To demonstrate these functions, we performed cross-modal analyses linking gene expression datasets with neuronal reconstructions. In larval zebrafish, we utilized previously published and registered transgenic labeling for the dopamine D2a receptor *drd2a*^12^ and whole-brain hybridization chain reaction (HCR) image for the dopamine D4a receptor *drd4a*^14^ (Figure 4B, I). For each loaded 3D image, iBrAVE can render expression intensities (Figure 4B, II-i) and convert above-threshold expression patterns into spatial feature points, which can be mapped to annotated brain regions (Figure 4B, II-ii) and allows generation of expression density maps (Figure 4B, II-iii). By spatially mapping each gene image onto the nervous system-wide single-neuron morphology database,^15^ we identified the corresponding neuronal populations (Figure 4B, II-iv; see Methods). By performing HCR-FISH on sparsely labeled fish samples used for single-neuron reconstruction (see Methods), we validated that, among the mapped *drd2a*+ and *drd4a*+ populations, two reconstructed mitral cells (MCs) with distinct projection patterns were confirmed to express *drd2a* and *drd4a*, respectively (Figure 4B, II-v). Boolean intersection of the two images (in Figure 4B, I) identified areas with overlapping expression of *drd2a* and *drd4a* (Figure 4B, III-i). These areas were then used to filter the *drd2a*+,*drd4a*+ neuronal population, which localized predominantly to the pineal gland (Figure 4B, III-ii). Consistently, previous studies reported the expression of both *drd2a* and *drd4a* in the pineal gland.^57,58^ Our whole-mount HCR-FISH results further showed the juxtaposed *drd2a*+ and *drd4a*+ puncta within the pineal gland (Figure 4B, III-iii), supporting their co-expression in pineal neurons.

In the mouse brain, we analyzed MERFISH-based spatial transcriptomics data,^22^ which were imported into iBrAVE as feature points (see Methods). Spatial distributions of cells expressing four subtypes of dopamine receptor genes (*Drd1* and *Drd5* representing D1-type receptor (D1R), and *Drd2* and *Drd3* representing D2-type receptor (D2R)) were compared, revealing distinct cell-resolved gene expression abundance (Figure 4C, I) and regional distribution densities (Figure 4C, II). Within the hippocampal formation (HPF; Figures 4D and 4E), quantitative analysis of cell density across subregions revealed a more enrichment of D1R-expressing cells compared with D2R-expressing cells, with *Drd5*-expressing cells showing particularly strong prevalence. This HPF bias contrasted with the whole brain, where the densities of D1R- and D2R-expressing cells were comparable (Figure 4E, left bar). By mapping the positions of gene-expressing cells onto soma locations from hippocampal single-neuron morphology reconstructions,^23^ iBrAVE reveals the morphologies of neurons corresponding to the expression domains of distinct dopamine receptors (Figure 4F). Sholl analysis revealed that D2R-mapped neurons exhibited significantly more intersections than D1R-mapped neurons, particularly at distances of ∼800 - 1,800 μm from the soma (Figure 4G). Further analysis of a subset of reconstructed neurons with identified dendritic and axonal arbors showed that this difference was mainly attributable to axonal rather than dendritic compartments (Figures S2A and S2B). Consistently, compared to D1R-mapped neurons, D2R-mapped neurons exhibited greater values across multiple morphological properties, including total length, bifurcation number, branch number, terminal number, and maximum branch order, indicating greater overall structural complexity (Figures 4H, and S2C–S2H).

These findings suggest that hippocampal D2R-mapped neurons possess more complex arbors than their D1R-mapped counterparts.

These analyses demonstrate that iBrAVE enables the identification of candidate neurons expressing specific genes, thereby facilitating the association of molecular markers with characteristic morphologies. Conversely, it allows searching for molecular markers corresponding to neurons with characteristic morphological features. These capabilities bridge genetic and morphological classification, advancing an integrative multimodal framework for neuronal typing.

### Cross-modal analysis: linking neuronal activity to circuit

Through spatial correspondence, iBrAVE can also integrate functional and structural datasets within a common 3D spatial framework, enabling whole-brain neuronal activity to be mapped onto underlying circuit architectures (Figure 5A). As a case study, we analyzed whole-brain calcium imaging data recorded from freely swimming larval zebrafish engaged in prey capture behavior.^59^ Neuronal activity dynamics during prey capture were registered to the zebrafish CPSv1 (see Methods) and overlaid with reconstructed single-neurons and connectivity datasets.^15^ This integration enables frame-by-frame linkage of neuronal activity with morphological reconstructions and brain regions, thereby associating distinct behavioral epochs—including prey detection, J-turn, approach, eye convergence (EC), inhale, swallow, and digestion—with specific morphological ensembles distributed in whole-brain regions (Figures 5B, 5C, and Video S4). Notably, corresponding neurons activated during individual epochs displayed both shared and divergent morphologies and projection patterns (Figure 5C), indicating a combination of continuous transitions and modular recruitment of circuit components across different behavioral stages.

Among corresponding neurons activated during prey detection, contralaterally descending projection neurons in the OTAOS (optic tract and accessory optic system) region were identified to directly receive RGC inputs (Figure 5D, I). These neurons belong to the glutamatergic morphotype #Glu-2C0203^15^ and exhibit similar soma location and projection pattern to the previously reported neuron class that can trigger predatory behavior.^60^ We named them pre-capture initiation neurons (PCINs). Using iBrAVE’s neuronal wiring calculation *Filter*, we traced circuit connections within the activated ensembles and revealed diverse neurons involved. During the initial detection epoch, in addition to the visual-related PCIN, we observed activation of olfactory pathways, including olfactory sensory neurons (OSNs) and their downstream MCs in the olfactory bulb (Figure 5D, II-i). These MCs were further connected to spinal cord-projecting, motor-related neurons in the dopaminergic diencephalic cluster 2 (DC2)^61^ (Figure 5D, II-ii) and to visual-related projection neurons in the optic tectum (OT) via subpallium (Sp) (Figure 5D, II-iii). These two putative olfactory-relevant circuits require further experimental verification.

During approach and EC epochs, the identified downstream partners of PCINs included motor coordination-related neurons, such as oculomotor neurons (nIV, the trochlear nerve motor nucleus), reticulospinal neurons (RSNs), and the nucleus of the medial longitudinal fasciculus (nMLF) neurons, which were relayed via an ascending hindbrain (H) neuron (Figure 5E, I). Furthermore, we identified long-range ascending neurons downstream of the nMLF and RSNs that projected feedback to multiple sensory regions, including known cholinergic neurons in the octavolateralis efferent nucleus (OEN)^62^ projecting to lateral line neuromasts (mechanosensory) (Figure 5E, II-i), and hindbrain neurons projecting to the olfactory bulb (olfactory) (Figure 5E, II-ii) or optic tectum (visual) (Figure 5E, II-iii). While future experimental validation is required, these candidate feedback pathways highlight potential cross-modal coordination during prey capture.

Altogether, these analyses illustrate that iBrAVE enables data-driven discovery of structure-function relationships at cellular resolution across behaviorally relevant circuits. By unifying multimodal datasets—including neuronal activity, morphology, and connectivity—within a common spatial reference, iBrAVE provides a platform for *in silico* hypothesis generation.

### Cross-scale analysis: integrative LM-EM neuron annotation

Microscale EM datasets often lack cell-type identity, whereas mesoscale LM datasets typically lack synaptic connectivity information. To leverage their complementary strengths, we developed a cross-scale LM-EM matching function within iBrAVE (see Methods). It aligns LM and EM datasets based on single-neuronal soma position and morphological similarity, thereby (1) transferring LM-based structural and functional cell-type labels to EM reconstructions, and (2) mapping EM-derived synaptic connectivity onto LM reconstructions.

For EM reconstructions, iBrAVE supports joint visualization of EM-derived neuronal morphologies and their synaptic connections (Figure 6A, I), allowing dynamic querying of upstream and downstream partners based on annotated synaptic relationships. For example, by interactively selecting a single postsynaptic site on a locus coeruleus (LC) neuron, an individual presynaptic neuron can be retrieved and visualized (Figure 6A, II). To infer the cell-type identity of this upstream partner, we performed morphology-based matching against LM reconstructions in the ZExplorer single-neuron morphology atlas.^15^ This analysis revealed three neurons in close proximity with high similarity scores, all belonging to the glutamatergic morphotype #Glu-5L0701^15^ (Figure 6B). Neurons of this morphotype correspond to *vglut2a/2b*-positive dopaminergic neurons in DC6,^61,63^ suggesting that the LC may directly receive dopaminergic inputs.

Conversely, LM neurons can be linked to EM reconstructions via morphology-based matching, enabling detailed interrogation of their synaptic connectivity. For example, within the corresponding neuronal ensemble that was active during the detection epoch of prey capture behavior, we identified a neuron whose location and morphology correspond to the spinal cord-projecting A11-type dopaminergic neurons in DC2^61,64^ (see Figure 5D, II-ii). Matching this neuron to EM reconstructions yielded a high-similarity candidate located in DC2 (Figure 6C, I). This candidate was labeled with cytosolic enhanced ascorbate peroxidase-2, a marker of monoaminergic neurons in the EM data,^16^ confirming its dopaminergic identity. Further inspection of the EM connectome revealed a presynaptic partner in DC5 (Figure 6C, II). Based on its soma location and morphology,^61^ this partner is likely a dopaminergic neuron as well, suggesting interconnections between dopaminergic nuclei.

This bidirectional LM-EM matching establishes an iterative closed-loop, whereby LM motivates synapse-level EM exploration, and EM-derived connected neurons are mapped back to LM to refine cell-type annotations. Together with the cross-modal analyses, this integration of LM’s cellular-resolution reconstructions with EM’s synapse-level details provides a unified framework for cross-scale, cross-modal mapping of synaptic connectivity, neuronal identity, and neuronal activity.

## DISCUSSION

iBrAVE addresses a significant challenge in current brain atlas research: the lack of a unified solution for integrative analysis of 3D brain atlas datasets across modalities and scales in diverse species. By abstracting diverse datasets into a set of generalizable data types—volumetric images, parcellated regions, spatial feature points, neuronal tracings, and networks—iBrAVE accommodates existing brain atlas data, from gene expression to neuronal morphology to neuronal activity, independent of species. It enables researchers to conduct a continuous analysis workflow, spanning image registration, modality-specific quantification, and cross-modal mapping and integrative interrogation. This capability underpins its importance in providing a foundational solution for integrative deciphering of the molecular, structural, and functional architecture of the brain.^4,65^

One of the key innovations of iBrAVE is its capability to support 3D interoperability of multimodal brain data, in accordance with the FAIR guiding principles.^66^ By registering diverse datasets into a species-specific anatomical coordinate framework, iBrAVE enables dynamic crosstalk between any two modalities based on spatial relationships, for example, mapping gene expression onto neuronal morphologies, mapping brain-wide activity patterns to structural connectivity, or matching LM and EM reconstructions. Ultimately, this framework enables holistic integration of diverse modalities, providing comprehensive multimodal annotations for a single biological entity. In addition, iBrAVE couples quantitative analysis with interactive visualization, ensuring that computational results are transparent and intuitively interpretable. Analyses are thus directly anchored to visual evidence, promoting an iterative “see to know, and know to see” cycle of discovery.

Another distinctive feature is its high accessibility. iBrAVE’s intuitive, GUI-based interface, featuring context-sensitive right-click menu interactions directly within the 3D rendering window, removes barriers for users lacking programming expertise. This contrasts with code-intensive toolkits, significantly broadening potential application within the neuroscience community. Furthermore, its built-in support for saving, restoring, and recording analysis workflows ensures reproducibility and fosters collaborative development among researchers.

Technically, iBrAVE leverages a distributed computing architecture to support high-performance 3D rendering and real-time interaction, even with large-scale datasets such as whole-brain neuronal reconstructions or synaptic connectomes. It also supports algorithmic extensibility, allowing researchers to incorporate custom analytical methods (such as neuronal classification algorithm) while taking advantage of iBrAVE’s visualization and data mining capabilities. Multiple export options, ranging from structured tables and high-resolution figures to animations and immersive VR content (Video S5), along with real-time 3D interactive visualization capabilities, further expand its utility for scientific communication, teaching, and public engagement.^67^

While initially developed for neuroscience research, iBrAVE’s design principles—spatially anchoring of data, standardized formats for common biological modalities, and cross-modal integration—make it applicable across biological systems. Any system in which molecular, structural, and functional datasets can be registered to a common anatomical framework can benefit.^68^ For example, iBrAVE can be extended to atlas-based investigations of the heart, lung, or kidney, across developmental or pathological states. This generality positions iBrAVE as a versatile spatiotemporal platform for systems biology beyond the brain.

We envision iBrAVE operating within a dual-system ecosystem. The standalone platform functions as an exploratory engine for cross-modal and cross-scale visual analytics, whereas a web-based interface serves as a data query and structured knowledge portal powered by graph-based ontology^69,70^. Altogether, they form a mutually reinforcing loop: findings from iBrAVE feed into the ontology, while ontology-guided exploration drives targeted analyses within iBrAVE. This interplay promotes both hypothesis-driven and data-driven discovery, while supporting open science practices and collaborative exchanges.

In summary, iBrAVE provides not only a technical solution but also a conceptual framework for integrative neuroscience research. We expect iBrAVE to catalyze a paradigm shift in experimental design, in which spatial registration to standardized coordinate systems becomes a guiding principle for data acquisition. Such spatial anchoring not only enhances comparability but also incentivizes harmonization of datasets across laboratories. By unifying multimodal data into a spatially coherent and accessible environment, iBrAVE can accelerate the emergence of community-driven, continuously updated atlases that increase in scope and resolution over time, fostering a shared and cumulative scientific resource.

## Limitations of the study

The integrative analysis of multimodal, multiscale data in iBrAVE is built upon spatial correspondence among datasets aligned within a common reference space. Although iBrAVE allows user-defined thresholds for associating features across modalities, the fidelity of cross-modal mapping is inherently constrained by the quality of input data and the accuracy of registration. Future developments are needed to explicitly quantify or visualize the uncertainty associated with these spatial alignments. In addition, experimentally established ground-truth correspondences among multiple modalities at the single-cell level—such as jointly measured molecular, morphological, and functional properties—are not yet natively supported for direct import and comparison. Supporting such ground-truth multimodal relationships would enable systematic validation, benchmarking, and integration with mappings inferred from spatial proximity. More broadly, while iBrAVE accommodates user-defined analytical approaches, as exemplified by clustering of single-neuron morphology, such extensions currently require direct integration of algorithms into the analysis and visualization codebase rather than invocation through a unified interface. Future work will develop a generic plugin architecture for analytical modules, allowing new computational methods to be incorporated as data acquisition and analysis methodologies continue to advance. By combining extensible analytics with iBrAVE’s visualization and data-mining capabilities, the framework enables community-driven development and continued expansion of its application scope.

## RESOUCE AVAILABLITY

### Materials availability

Comprehensive documentation, including video- and text-based tutorials, detailed algorithmic descriptions, and developer resources, is available at: https://zebrafish.cn/iBrAVE.

### Data and code availability

The multimodal datasets across species used in this study are publicly available (see Key Resources Table and Table S1). iBrAVE (built on Paraview 5.12.0) is fully open source. All source code and software packages are available at https://github.com/soaringdu/Proj-iBrAVE as of the publication date.

## ACKNOWLEDGEMENTS

We are grateful to Dr. Zhifeng Yue for selecting the ParaView infrastructure and contributing to the framework design. We thank Dr. Chun Xu, Yu Mu, Shengjin Xu, and Tielin Zhang from the advisory committee of the institutional Center for Data and Computing in Brain Science; Tingting Zhao from the Zebrafish Group of the institutional research facility for Mapping Brain-Wide Mesoscale Connectome; and Jun Zhang from Achievement Transformation Office for their support in the implementation and documentation of iBrAVE. We thank Tianlun Chen, Qimeng Zhao, Meiyu Zheng, Xin Wang, Hongli Wan, and Tingting Zhao for their assistance with software tutorials. We thank Hao Yang from Dr. Zhifeng Liang’s laboratory for assistance with macaque transcriptomic data processing, Dr. Chao Li for assistance with HCR-FISH experiments, and Shan Wang for help with VR-related video recording. We thank Dr. Misha B Ahrens for providing the *Tg(elavl3:H2B-GCaMP6f)* fish line. This work was supported by Brain Science and Brain-like Intelligence Technology - National Science and Technology Major Project (2021ZD0204500, 2021ZD0204502, 2021ZD0204504, and 2022ZD0209600), the National Natural Science Foundation of China (32321003, 62320106010, 32400817, 62276253, and U22B2063), Shanghai Municipal Science and Technology Major Project (2025SHZDZX025D03 and 25511102500), Shanghai Municipal Commission of Economy and Informatization (2025-GZL-RGZN-BTBX-02005), and Key Research Program of Frontier Sciences (QYZDYSSW-SMC028) and Strategic Priority Research Program (XDB32010200) of Chinese Academy of Sciences. X.F.D. is also supported by the Youth Innovation Promotion Association, Chinese Academy of Sciences.

## AUTHOR CONTRIBUTIONS

X.F.D. and J.L.D. conceived and supervised the study. X.F.D., J.L.D., and Y.F.W. designed the overall framework. J.F.W. developed the software with input from H.B.M., with Y.W.Z. contributing to quality control. M.Q.C., H.B.M., J.F.W., Y.W.Z., S.X.C., and J.Q.G. prepared the software tutorials with input from X.F.D. and Y.F.W.. C.X.J. designed the software logo, boot animation, and icons. F.N.L. provided the EM data. T.L.Z acquired brain functional imaging data from freely moving zebrafish during prey capture behavior, under the supervision of K.W.. C.C. and Y.F.W. conducted HCR-FISH experiments under the supervision of S.J.X.. X.D.Z performed confocal imaging. Y.M.Z. processed the spatial transcriptomics data. Y.F.W. and M.Q.C. analyzed the data and prepared illustrations with input from H.B.M. and X.F.D.. Y.F.W. and M.Q.C. generated the videos. X.F.D. and J.L.D. wrote the manuscript with input from all authors.

## METHODS

### Zebrafish

Adult zebrafish were maintained at 28 ℃ in an automatic housing system at the National Zebrafish Resources of China (Shanghai, China). Experimental larvae were raised in 10% Hanks’ solution containing (in mM): 140 NaCl, 5.4 KCl, 0.25 Na_2_HPO_4_, 0.44 KH_2_PO_4_, 1.3 CaCl_2_, 1.0 MgSO_4_ and 4.2 NaHCO_3_ (pH 7.2) under a 14 h:10h light:dark cycle at 28 ℃. The following transgenic line was used for calcium imaging: *Tg(elavl3:H2B-GCaMP6f)jf7*.^71^ Larvae sparsely labeled with tdTomato in neurons at 6 days post-fertilization (dpf), used for multiplexed fluorescent *in situ* hybridization (FISH), were derived from preserved samples previously prepared for single-neuron morphology acquisition.^14^ All animal procedures were approved by the Animal Care and Use Committee of the Center for Excellence in Brain Science and Intelligence Technology, Chinese Academy of Sciences (NA-046-2023).

### iBrAVE layout

Upon startup, iBrAVE presents an initialization interface promoting users to select a species-specific reference brain, which establishes the coordinate framework for subsequent analyses. Users can also define the operating mode as either visualization-only or visualization-and-analysis. Once launched, iBrAVE displays a modular interface comprising seven core components:

1. Menu Bar – Provides essential functionalities, including data import/export, 3D view configurations, volumetric data cropping and slicing, custom viewpoint management, animation playback, and access to the *Filter* toolbar. The *Filter* toolbar enables one-click access to built-in analytical modules, such as morphological quantification, neuronal wiring calculation, and network connectivity analysis.
2. Pipeline Browser – Organizes all imported datasets and analytical outputs hierarchically, allowing users to track and manage data processing workflows.
3. Data Browser – Lists all loaded neuronal tracings, brain regions, image volumes, and feature points in structured modules. It supports searching, sorting, toggling visibility, customizing colors, and interactive highlighting.
4. Properties Panel – Enables parameter adjustment for items in the processing pipeline, such as color, opacity, rendering settings, and analytical thresholds.
5. 3D RenderView – Serves as the central environment for interactive 3D visualization, facilitating real-time exploration of multimodal brain data and derived analytical results.
6. Context-Sensitive Right-Click Menu – Offers shortcut operations within the RenderView, including toggling visibility of all or specific data types, switching color-coding schemes of neuronal tracings, executing group-based operations (see the section “Group function” in Methods), exploring neuron-region relationships and inter-neuronal connectivity, and exporting projection or connectivity matrices.
7. Animation View – Allows users to define camera trajectories, control display sequences, and adjust rendering settings to generate customized animations of multimodal brain data.

### Basic tutorials

We illustrate iBrAVE’s workflow using neuron-centric visualization and analysis as an example, demonstrated with the zebrafish CPSv1 reference atlas:

1. Launching software and selecting reference atlas

Select “Zebrafish CPSv1” in the startup interface and click “Start”.

2. Loading multimodal data

Click “Open” to import datasets. In the dialog box, select folders containing neuronal tracings and brain regions. Assign appropriate readers (e.g., Neuronal Tracing Data Folder Reader or Region Data Folder Reader) and click “Apply”.

3. Visualizing data and controlling visibility

Toggle dataset visibility using the eye icons in the Pipeline Browser or Data Browser. Within the RenderView, right-click to open the context menu and select “Show Only Neuronal Type” → [specific type] to isolate neurons by type (see the section “Neuronal type annotation” in Methods), or “Show Only Neural Structure” → “Soma/Axon/Dendrite” to display specific compartments.

4. Analyzing neuronal morphology

Select a neuronal tracing data in the Pipeline Browser, click the “Neuronal Morphology” *Filter* icon in the *Manu Bar*, select the analysis target (entire neuron, axon, or dendrite) in the *Properties Panel*, and designate an output directory. Click “Apply” to run the analysis, and results are exported as Excel spreadsheets.

5. Exploring neuron-region relationships

Right-click a neuron to open the context menu and select “Show Only Neuron” → [regions with specified relationship] to isolate the neuron alongside its soma-locating, dendrite-innervating, or axon-projecting regions. Conversely, right-clicking a brain region and selecting “Show Only Region” → [neurons with specified relationship] retrieves neurons associated with that region.

6. Exporting projection metrics

Within the *RenderView*, right-click to open the context menu and navigate to “Analysis And Export” → “Projection Weight” → “Neuron-Region” → [specific projection metric]. Specify a save location; results are exported as structured Excel files for downstream analyses.

7. Managing sessions

Save or restore sessions via “File” → “Save State…” and “File” → “Load State…” for continuity or reproducibility.

### Neuronal type annotation

iBrAVE supports visualization and analysis of imported neuronal tracing data by user-defined neuronal types, organized in a two-tier hierarchy (main type and subtype). Type-based operations, such as toggling visibility and assigning colors, can be quickly accessed via the context menu or “Data Browser”. Projection-related analyses, including projection heatmap visualization and projection weight matrix export, can be automatically executed for each neuronal type.

Two annotation methods are supported: (1) Filename-based annotation, where the subtype name is included in the filename, prefixed by an underscore (“_”), and mapped to a main type in the source code (e.g. subType2MainTypeMap[“vglut2a”] = “Excitatory”); and (2) JSON-based annotation, where hierarchical type information is embedded directly in the neuron JSON file (see the section “JSON file applications” in Methods). When detected, type information is shown in the context menu and displayed via tooltips upon hover.

### Group function

iBrAVE enables selective management of subsets of imported data through a “Group” function, supporting targeted visualization and analysis of multimodal elements such as neurons, brain regions, and feature points. Groups can be defined through two methods: (1) Box selection, activated via the context menu item “Select into Group”, enabling rectangular region selection on screen; and (2) Pick selection, allowing users to right-click individual elements, view metadata, and add/remove them from groups via “Add into/Remove from Group” → “Picked” menu items. Group states can be saved/loaded via “Save/Load Group State” menu items for reproducible workflows.

Grouped data can be manipulated like general data, including exporting basic information, toggling visibility, adjusting global rendering settings (e.g., color, opacity), and exploring neuron-region relationship and neuronal connectivity. iBrAVE also provides a dedicated panel for creating and managing multiple groups, enabling combinatorial visualization and comparative analysis within a single RenderView.

### Neuronal wiring analysis based on morphology

Neuronal connectivity was inferred using the Neuronal Wiring Calculation *Filter*, which evaluates axodendritic spatial proximity in single-neuron reconstructions. Four computational models were implemented: center, cubic, cylinder, and tube. The first three assess the local proximity of individual neurite compartments, while the tube model evaluates entire axonal or dendritic volumes. Users can define proximity thresholds based on model radii (see Figure 3E, top) and further filter connectivity using either the number of qualified compartment pairs (center model) or volumetric overlap (absolute or relative; cubic, cylinder, and tube models). Analyses can be refined by restricting branch types, such as fork-fork or fork-terminal branches. The resulting directed and weighted connectivity networks can be exported, with nodes representing individual neurons or neuronal clusters and edge weights defined by either compartment pair counts or overlapping volume.

### Cross-modal mapping based on spatial queries

To enable cross-modal mapping, iBrAVE provides a spatial data filtering framework that links data with structural, molecular, or functional features. Take neuron filtering as an example, filtering can be performed based on images or feature points. In the image-based mode, soma positions are evaluated against volumetric images (e.g., immunohistochemistry, calcium imaging), and neurons are retained if the corresponding voxel intensity falls within a user-defined range. In the feature point-based mode, Euclidean distances between neuron somata and feature points (e.g., gene-expressing cells, multi-unit recording sites) are calculated, and neurons are retained if they lie within a user-defined proximity threshold, optionally constrained by feature-specific scalar values (e.g. gene expression level, neural activity strength). The output includes the resulting neuron subsets and a point cloud of the qualifying image voxel locations or spatial feature points for subsequent visualization and quantitative analysis.

### LM-EM matching based on morphological similarity

To enable correspondence between LM and EM reconstructions, iBrAVE provides a Neuron Similarity *Filter*. Neurons are represented as SWC point clouds, and pairwise similarity is computed using distance-based metrics, including Chamfer (the bidirectional average closest point distance), directed Chamfer (forward and reverse), centroid Euclidean, and Hausdorff distances. Distances are converted to similarity scores as inverse functions of (1 + distance). To improve efficiency, candidate pairs are first pre-filtered by soma proximity (user-defined SomaDistanceThreshold), followed by fast nearest-neighbor searches using k-d trees and parallel processing of neuron pairs. For each neuron, similarity scores are ranked, and only the top N partners (specified by Neuron Count in the *Properties Panel*) above a user-defined similarity threshold are retained. Comparisons can be restricted within or between neuronal clusters (ClusterRelation). The output provides ranked similarity lists for each neuron, which can be exported in CSV/JSON formats or directly visualized in iBrAVE, thereby enabling systematic matching of LM morphotypes with EM-derived reconstructions.

### JSON file applications

iBrAVE employs JSON (JavaScript Object Notation) files to store metadata for multimodal datasets. Metadata is organized through predefined keywords written as associative arrays in JSON format. Each dataset entry is indexed by a datum ID, which acts as a unique identifier linking to the value lists of various relationship-specific keywords. For example, in a neuron JSON file, the keywords “type_array”, “output_region_array”, and “upstream_neuron_array” specify the hierarchical neuronal type, IDs of target projection regions, and IDs of potential input neurons (see the section “Neuronal wiring analysis based on morphology” in Methods), respectively.

Users can create JSON files manually or export them directly from iBrAVE after data processing. To export, right-click in the RenderView layout to open the context menu, and navigate to “Analysis and Export” → “General Information” → “All in Separate/Single JSON file”. To apply JSON files, place them in the corresponding data folder and name them info.json. When loading data, iBrAVE automatically detects info.json and prioritizes it as the source of input metadata. This eliminates redundant computation, which is particularly beneficial for resource-intensive tasks such as neuron projection analysis and neuronal/synaptic wiring analysis in large-scale datasets.

### Performance benchmarking

We benchmarked iBrAVE in standalone and client-server modes. Standalone tests were conducted on a DELL Precision T7920 Tower workstation (Intel® Xeon® Gold 6250 CPU, 3.90□GHz; 768□GB RAM; dual 1.86□TB NVMe SSDs; NVIDIA GeForce RTX 3060 GPU, 12□GB). Client-server tests used the same workstation as the client and a high-performance computing (HPC) cluster as the server, coordinated by SLURM. HPC nodes were configured with dual AMD EPYC 7742 64-core processors (2.30□GHz) and dual NVIDIA A100 GPUs (40□GB each), connected to the storage system via InfiniBand. Server instances were launched with MPI using mpirun -np 4 pvserver.

Three neuron reconstruction datasets were tested: (i) 5,000 zebrafish neurons (264□MB; 14,153,576 nodes), (ii) 5,000 mouse neurons (12,000□MB; 458,603,220 nodes), and (iii) 2,000 macaque neurons (887□MB; 30,547,138 nodes). Prior to each test, the system was reinitialized to ensure consistent runtime environments. Each test was repeated six times under both deployment modes, with execution log files captured (-l=log.txt,9) to record runtimes. Execution times (in seconds) were extracted from log files for performance evaluation.

### Preprocessing of spatial transcriptomics data of mice and macaque

MERFISH data (MERFISH-C57BL6J-638850) from an adult C57BL/6J mouse brain were obtained from the Allen Brain Atlas (https://alleninstitute.github.io/abc_atlas_access/descriptions/WMB_dataset.html). For Figure 2, cell coordinates pre-mapped to the Common Coordinate Framework (CCFv3) were integrated with cell-type annotations. One single output file (columns: cell-type identifier, *x*, *y*, *z*) was generated for iBrAVE visualization. For Figure 4, raw gene counts were processed in R (v4.2.2) with Seurat (v5.0.1), normalized with SCTransform using variance-stabilizing transformation (clip range [–10, 10]), and integrated with cell coordinates. For each gene, final outputs (columns: cell-type identifier, *x*, *y*, *z*, normalized gene expression) were exported for iBrAVE visualization and analysis. Stero-seq data from macaque brain slices (“macaque1”) were obtained from https://macaque.digital-brain.cn. Cell coordinates and cell-type annotations were processed analogously.

### Digitization of zebrafish gene expression atlas

The mapzebrain volumetric image atlas of cellular-resolution gene expression^14^ was registered to the zebrafish CPSv1 using an inter-atlas transformation.^15^ Cell positions (soma centroids) expressing specific genes were manually annotated in Fiji using the Cell Counter plugin and then transformed to the CPS using antsApplyTransformsToPoints in ANTs.^72^ For each gene, a feature-point CSV file containing cell coordinates was generated for iBrAVE visualization and analysis.

### Multiplexed hybridization chain reaction-fluorescence *in situ* hybridization (HCR-FISH)

Larvae with sparsely tdTomato-labeled neurons were fixed in 4% paraformaldehyde (PFA) at 4 °C for 24 h, washed twice with 1× phosphate-buffered saline (PBS), and transferred to 100% methanol at 4 °C for long-term storage. Prior to hybridization, samples were rehydrated by two 10-min washes in 2× saline–sodium citrate (SSC) buffer at room temperature and photobleached overnight at 4 ℃ using an 18 W white LED to minimize tissue autofluorescence.

HCR-FISH was performed following a modified protocol.^73^ Briefly, samples were post-fixed, permeabilized, and digested with proteinase K before hybridization with custom-designed DNA probes (synthesized by Beijing Tsingke BioTech) targeting *drd2a* (ID: SHP25061100009), *drd4a* (ID: SHP25061100010), and *tdTomato* (ID: SHP25011000003). After stringent washes, signal amplification was achieved using snap-cooled fluorescent hairpins conjugated to Alexa Fluor 488, Alexa Fluor 546, or JF-669 (synthesized by Sangon Biotech) in amplification buffer (Molecular Instruments). Excess hairpins were removed with a 5× SSCT wash, and samples were stored at 4 ℃ in the dark until imaging.

### Confocal imaging

Samples were embedded in 2% low-melting-point agarose in 1× PBS and imaged with an Olympus FV3000 upright confocal microscope with a 20× water-immersion objective (N.A. 1.0). *Z*-stacks were acquired in the upward direction at a voxel resolution of ∼0.49 × 0.49 × 2.5 μm. To compensate for fluorescence attenuation, laser power was progressively increased along the *z*-axis. Excitation wavelengths were 488 nm, 561 nm, and 640 nm for Alexa Fluor 488, Alexa Fluor 546, and JF-669, respectively.

### Zebrafish prey capture behavior

Transgenic larvae *Tg(elavl3:H2B-GCaMP6f)jf7*^71^ expressing a pan-neuronal calcium indicator in a *nacre* background, were used for behavioral imaging. Larvae were raised at 28 °C and imaged at 6 dpf. To enhance prey-capture motivation, individual larvae were food-deprived for 12 h before recording. Each larva was placed in a custom imaging chamber (45 mm diameter, 0.8 mm depth) containing 10% Hanks’ solution and ∼200 paramecia as prey. The chamber was covered with a 1 mm-thick coverslip to ensure a stable imaging environment.

### 3D#tracking for whole-brain neuronal Ca^2+^ imaging in freely swimming zebrafish

Real-time 3D tracking was implemented similarly to a previous demonstration^59^ to enable stable whole-brain calcium imaging in freely swimming larvae. Briefly, larvae were illuminated with near-infrared (NIR) LEDs, and a high-speed camera (Basler ace acA1920-155μm; 1.5□ms exposure, 167□fps) captured scattered light for in-plane tracking. A custom LabVIEW program extracted fish centroid and orientation from each frame, and a control algorithm drove a two-axis direct-drive stage (Shanghai YIDING DDHD-4D54D5AP) to compensate for in-plane movement.

Axial (*z*-axis) tracking was achieved by a custom detector comprising a microlens array conjugated to the back pupil of the imaging objective and a camera (Basler aca2000–340km NIR; 90□fps) positioned at its focal plane. Depth was estimated by computing distance variations between images from individual microlenses. The positional error between the fish head and the set point was fed into a proportional-integral-derivative controller, which actuated a pair of piezoelectric stages (Harbin Core Tomorrow Science & Technology, P06.X500R) to stabilize axial position.

### Registration of brain-wide Ca^2+^ imaging data from freely moving zebrafish

For each fish, a reference brain was generated from a single 3D whole-brain imaging volume (600 × 600 × 250 voxels) with minimal motion artifacts. Principal axes were computed by covariance matrix analysis, and affine transformations were applied to standardize brain orientation and scale (resampled to 240 □×□500 □× □170 voxels). Other volumes were aligned to this reference via covariance-based axis alignment, cross-correlation-based translation optimization, and iterative refinement of rotation angles using maximum-intensity projections. Finally, aligned brain stacks of individual fish were registered to the zebrafish CPSv1 using ANTs.^15^

### Calcium imaging data analysis

Calcium imaging data were processed using a custom script to compute fluorescence changes (ΔF/F_0_). A brain mask was first applied to exclude non-brain regions. The baseline fluorescence (F_0_) was defined as the average intensity of six baseline frames acquired prior to the onset of prey capture behavior. For each frame, ΔF/F_0_ was calculated as (F−F_0_)/F_0_, scaled by a factor of 20 to enhance visual contrast, and converted to 8-bit format. The processed data were exported as multi-page TIFF files for direct import into iBrAVE.

### Statistical analysis

Plots and statistical analyses were performed using GraphPad Prism 9 (GraphPad Software) and MATLAB (Mathworks). Data are presented as mean ± SEM. For performance benchmarking of iBrAVE, group comparisons were conducted using multiple unpaired t tests with false discovery rate (FDR) correction for multiple comparisons (Figure S1). For Sholl analysis, differences between groups at each radius were assessed using the Wilcoxon rank-sum test with FDR correction for multiple comparisons (Figures 4G, S2A, and S2B). For morphological parameter analyses, between-group differences were assessed using the Wilcoxon rank-sum test (Figures 4H, and S2C–S2H). Sample sizes are indicated in the corresponding figure and/or figure legends. Statistical significance was defined as *P* < 0.05. **\****P* < 0.05; **\*\****P* < 0.01; **\*\*\****P* < 0.001.

---

**Figure S1.**
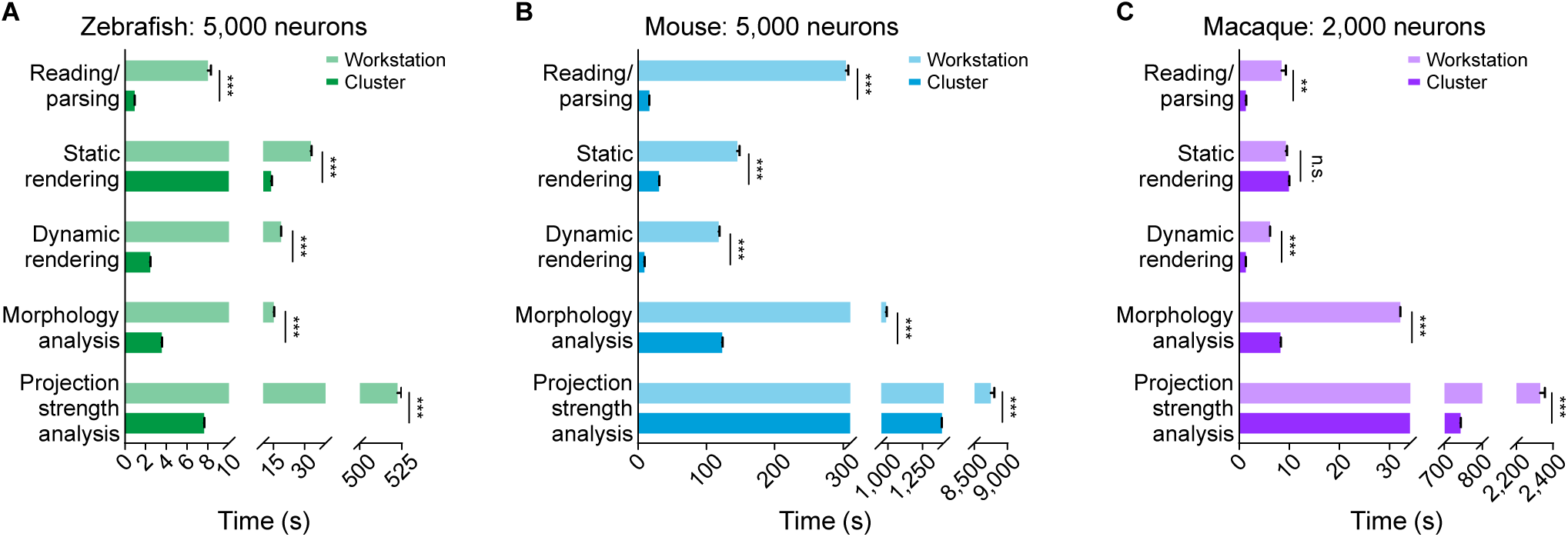

**Figure S2.**
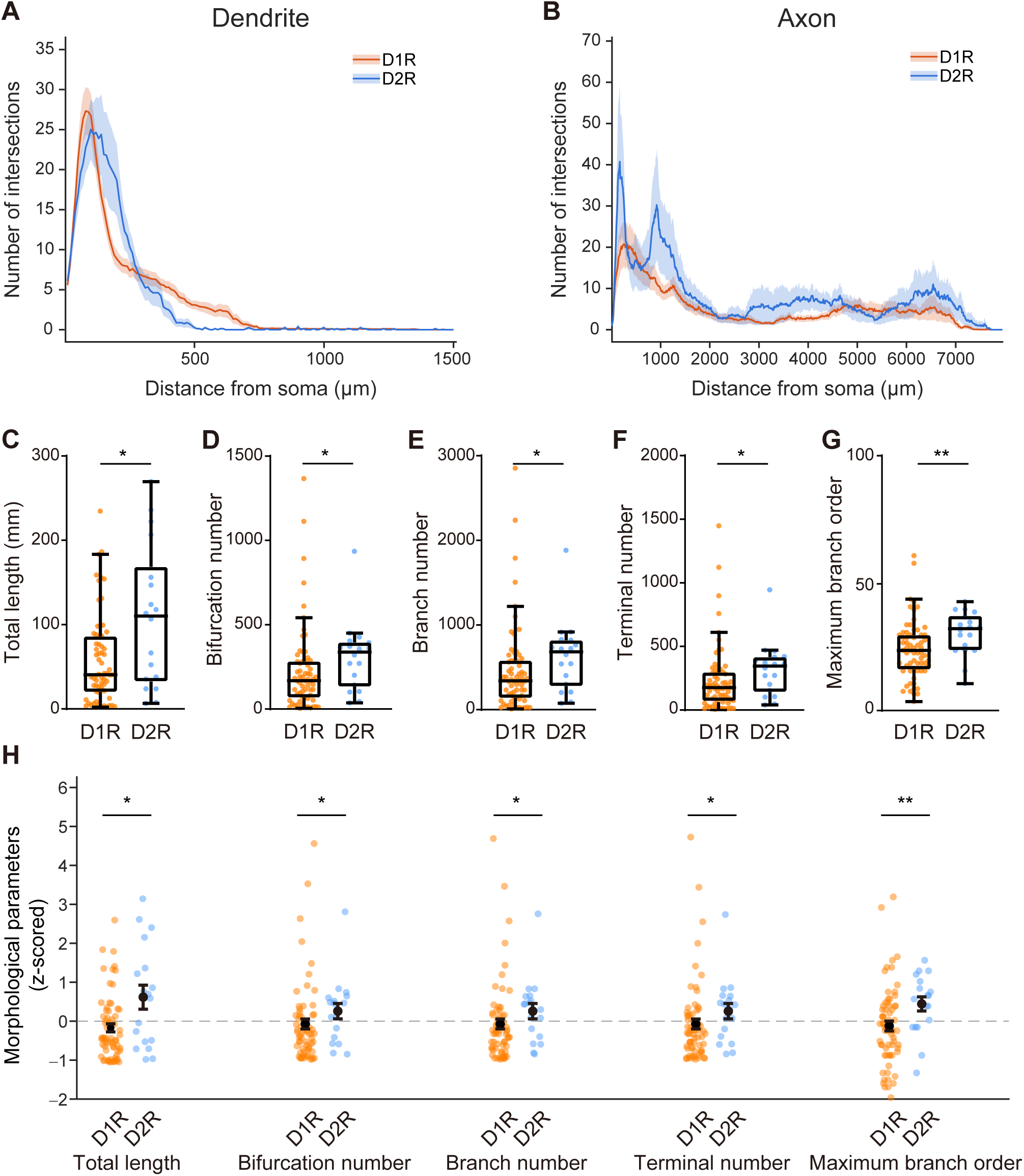

