## Supplementary information for "iBrAVE: a unified framework for 3D interactive and integrative analysis of brain atlas data across modalities and scales"

**Contents:**

Figures S1 and S2

Videos S1–S5

Tables S1 and S2

**Supplemental Figure Legends**

**Figure S1.** **Benchmarking iBrAVE performance across operations and deployment modes, related to Figure 2**

(A–C) Processing time for five operations—reading/parsing, static rendering, dynamic rendering, morphology analysis, and projection strength analysis—under standalone workstation (light) and computing cluster (dark) deployment modes, using single-neuron morphology reconstruction data from zebrafish (A, n = 5,000), mouse (B, n = 5,000), and macaque (C, n = 2,000). Bars show means from six independent runs, and error bars represent mean ± SEM. n.s., not significant; ***P* < 0.01, ****P* < 0.001 (Multiple unpaired t tests with false discovery rate correction).

**Figure S2. Morphological analyses of D1R- and D2R-mapped neurons in the mouse hippocampus, related to Figure 4.**

(A and B) 3D Sholl analysis of dendrites (A) and axons (B) in neurons exclusively mapped to D1R (n = 27) or D2R (n = 8). Intersections were quantified at 10-μm radial intervals from the soma center using concentric spheres. Solid lines represent mean values, and shaded areas indicate SEM.

(C–G) Box-and-whisker plots (Tukey style) with overlaid individual data points showing morphological parameters of neurons exclusively mapped to D1R (n = 65) or D2R (n = 18). Parameters include total length (C), bifurcation number (D), branch number (E), terminal number (F), and maximum branch order (G). ******P* < 0.05, *******P* < 0.01 (Wilcoxon rank-sum test).

(H) Z-scored morphological parameters from (C–G) for neurons exclusively mapped to D1R or D2R. Small dots represent individual neurons, and black symbols indicate mean ± SEM. The horizontal dashed line marks z = 0. **P* < 0.05, ***P* < 0.01 (Wilcoxon rank-sum test).

**Supplemental Videos**

**Video S1. Visualization of multimodal brain atlas datasets within the zebrafish common physical space, related to Figures 1 and 2**

**Video S2. Analytical navigation between neurons and brain regions, and visualization of projection strengths, related to Figure 3**

**Video S3. Interactive analysis and display of neuronal connectivity, related to Figure 3**

**Video S4. Real-time cross-modal visualization of whole-brain neuronal activities and underlying neural circuits during prey capture, related to Figure 5**

**Video S5. Immersive exploration of multimodal brain atlas data in virtual reality, related to Figure 1**

**Supplemental Tables**

**Table S1. The format, element count, size, vertex count, and source of the rendered atlas datasets across species, related to Figure 2**

| Data | Format | Elements | Size | Vertices | References | Source |
| --- | --- | --- | --- | --- | --- | --- |
| *Drosophila* | | | | | | |
| Brain template  (JRC 2018) | .obj | 46 | 15.6 MB | 252,470 | Bogovic et al., 2020^30^ | http://www.virtualflybrain.org |
| Gene expression  images | .nrrd | 7 | 102 MB | 834,159,480 | Jenett et al., 2012^50^ | http://www.virtualflybrain.org |
| Single-neuron  morphology atlas | .swc | 15,954 | 393 MB | 14,157,400 | Chiang et al., 2011^51^ | http://www.virtualflybrain.org |
| Brain activity  (Ca^2+^ imaging) | .nrrd | 1 | 400 KB | 409,600 | Mann et al., 2017^53^ | https://data.mendeley.com/datasets/8b6nw2xxhn/1 |
| Zebrafish | | | | | | |
| Nervous system  template  (ZExplorer CPSv1) | .stl | 103 | 345 MB | 3,619,087 | Du et al., 2025^15^ | https://zebrafish.cn/LM-Atlas/EI |
| Digitized gene  expression image  atlas | .csv | 107 | 3.8 MB | 126,739 | Shainer et al., 2023^14^ | https://mapzebrain.org |
| Single-neuron  morphology atlas | .swc | 20,011 | 1,136.6 MB | 66,015,256 | Du et al., 2025^15^ | https://zebrafish.cn/LM-Atlas/EI |
| Brain activity  (Ca^2+^ imaging) | .nrrd | 1 | 462.5 MB | 462,450,728 | This paper | This paper |
| Mouse | | | | | | |
| Brain template  (Allen CCFv3) | .obj | 566 | 55.3 MB | 486,674 | Wang et al., 2020^32^ | http://atlas.brain-map.org |
| Digitized spatial  transcriptomics | .csv | 1 | 208.6 MB | 3,739,961 | Yao et al., 2023^22^ | https://knowledge.brain-map.org/abcatlas |
| Single-neuron  morphology atlas | .swc | 28,721 | 6.6 GB | 228,319,756 | Gao et al., 2022^20^;  Qiu et al., 2024^23^  Gao et al.,  2026^24^ | https://mouse.braindatacenter.cn |
| Brain activity  (fMRI) | .nrrd | 1 | 178.5 KB | 89,100 | Yu et al., 2023^54^ | https://doi.org/10.12412/BSDC.1668502646.20001 |
| Macaque | | | | | | |
| Brain template  (NMTv2) | .stl | 196 | 183 MB | 1,918,048 | Seidlitz et al., 2018^36^ | https://afni.nimh.nih.gov |
| Digitized spatial  transcriptomics | .csv | 1 | 693.6 MB | 30,975,704 | Chen et al., 2023^26^ | https://macaque.digital-brain.cn/spatial-omics |
| Single-neuron  morphology atlas | .swc | 2,231 | 965.1 MB | 34,137,909 | Gou et al., 2025^27^ | https://zenodo.org/records/15128612 |
| Brain activity  (fMRI) | .nrrd | 1 | 442.6 KB | 110,592 | Yao et al., 2023^55^ | https://doi.org/10.12412/BSDC.1685411291.30001 |

**Table S2. The selected gene expression data for *Drosophila* and zebrafish, related to Figure 2**

| Species | Gene expression data |
| --- | --- |
| *Drosophila* | R77B11, R77E10, R77H05, R78B08, R78E02, R78G08, R79B01 |
| Zebrafish | *adcyap1a*, *agrp*, *alcamb*, *amylin*, *ascl1a*, *ascl1b*, *asip2b*, *atf5a*, *atf5b*, *avp*, *barhl1b*, *bhlhe23*, *bmp16*, *BX088596.1*, *c1ql3b*, *CABZ01073265.*, *calb1*, *calb2a*, *calb2b*, *calca*, *cart2*, *cart3*, *ccka*, *cckb*, *cebpa*, *chata*, *chodl*, *chrna6*, *CR361551.1*, *crabp1a*, *crha*, *crhbp*, *dacha*, *dazap1*, *dbx1a*, *dmbx1a*, *drd4a*, *emx2*, *esrrb*, *fabp7a*, *foxb1a*, *gad1a*, *gal*, *gbx2*, *gch1*, *ghrh*, *gjd2b*, *gnat1*, *grm2b*, *gyg1b*, *itpr1b*, *mafaa*, *mafbb*, *nefma*, *neurod1*, *ngb*, *nhlh2*, *nkx6.2*, *npb*, *npy*, *npy2r*, *npy2rl*, *npy7r*, *npy8br*, *nr4a2a*, *nxph2b*, *olig3*, *opn4b*, *otpb*, *pcp4a*, *pcp4l1*, *pdyn*, *penkb*, *pnoca*, *pnocb*, *pomca*, *pth2*, *pvalb6*, *pyya*, *rgs4*, *satb1b*, *sema3aa*, *six3a*, *six3b*, *slc17a7b*, *slc6a3*, *sox10*, *sox2*, *sp5l*, *sp9*, *sst1.1*, *sst1.2*, *sst5*, *sst7*, *stc2a*, *tac1*, *tac3b*, *tbr1b*, *tbx20*, *tbx2b*, *tfap2b*, *tfap2e*, *trh*, *txn*, *uts1*, *vip*, *zic2a* |
